# Lipid-Mediated Modulation of mGluR2 Embedded in Micelle and Bilayer Environments: Insights from Molecular Dynamics

**DOI:** 10.1101/2025.04.06.647485

**Authors:** Ehsaneh Khodadadi, Shadi A badiee, Ehsan Khodadadi, Mahmoud Moradi

**Affiliations:** Department of Chemistry and Biochemistry, University of Arkansas, Fayetteville, AR 72701

**Keywords:** mGluR2, G protein-coupled receptors (GPCRs), Cholesteryl hemisuccinate (CHS), POPC bilayer, Detergent micelle, Molecular Dynamics (MD) Simulations, Conformational dynamics, Transmembrane helices, Active and inactive states, Neurotransmitter signaling

## Abstract

Metabotropic glutamate receptor 2 (mGluR2), a subclass C member of the G protein-coupled receptor (GPCR) superfamily, is essential for regulating neurotransmitter signaling and facilitating synaptic adaptability in the central nervous system. This receptor, like other GPCRs, is highly sensitive to its surrounding lipid environment, where specific lipid compositions can influence its stability, conformational dynamics, and function. In particular, cholesteryl hemisuccinate (CHS) plays a critical role in stabilizing mGluR2 and modulating its structural states within cellular membranes and micellar environments. However, the molecular basis for this lipid-mediated modulation remains largely unexplored. To investigate the effects of CHS and lipid composition on mGluR2, we employed all-atom molecular dynamics simulations of mGluR2 embedded in both detergent micelles (BLMNG and CHS) and a POPC lipid bilayer containing 0%, 10%, and 25% CHS. These simulations were conducted for both active and inactive states of the receptor. Our findings reveal that CHS concentration modulates mGluR2’s structural stability and conformational behavior, with a marked impact observed within transmembrane helices TM1, TM2, and TM3, which constitute the core of the receptor’s transmembrane domain. In micelle environments, mGluR2 displayed unique conformational dynamics influenced by CHS, underscoring the receptor’s sensitivity to its lipid surroundings. Notably, a CHS concentration of 10% elicited more pronounced conformational changes than either cholesterol-depleted (0%) or cholesterol-enriched (25%) systems, indicating an optimal CHS range for maintaining structural stability. Our study provides atomistic insights into how lipid composition and CHS concentration impact mGluR2’s conformational landscape in distinct micelle and bilayer environments. These findings advance our understanding of lipid-mediated modulation in GPCR function, highlighting potential avenues for receptor-targeted drug design, particularly in cases where lipid interactions play a significant role in therapeutic efficacy.

## Introduction

G protein-coupled receptors (GPCRs) represent a large and diverse family of membrane-embedded proteins involved in transmitting external stimuli into intracellular responses. They play fundamental roles in mediating a broad spectrum of physiological processes by relaying signals across the cell membrane.^1–3^ Among the GPCR subtypes, metabotropic glutamate receptors (mGluRs) are particularly important for regulating synaptic activity and modulating neuronal responsiveness in the central nervous system. ^4^ mGluR2, a subclass C GPCR, contributes significantly to neurotransmitter signaling, impacting pathways responsible for synaptic strength, neuronal adaptability, and intercellular communication.^5,6^ The physiological relevance of mGluR2 is underscored by its role in supporting neural health, where perturbations in its signaling cascade have been implicated in a spectrum of neuropsychiatric and neurological conditions, such as schizophrenia, anxiety disorders, and depression.^7–10^ Deciphering the structural behavior and modulation of mGluR2 thus holds significant promise for therapeutic development.^11^

From a structural perspective, mGluR2 features a distinctive domain organization characteristic of class C GPCRs. It comprises an extracellular Venus Flytrap (VFT) module that mediates ligand recognition, linked to a seven-helix transmembrane domain (7TM) responsible for initiating intracellular signal transduction.^12^ Ligand binding induces conformational changes that propagate through both VFT and 7TM regions, initiating a downstream cascade of intracellular responses that modulate neuronal signaling and synaptic plasticity. ^13^ This signal propagation process is highly dependent on the surrounding lipid environment, as lipid-protein interactions influence receptor conformational states and downstream signaling efficacy.^14,15^ As with many GPCRs, the functional performance and structural stability of mGluR2 are tightly regulated by the composition and dynamics of the surrounding membrane lipids, particularly within the transmembrane domain.^16^

Among various membrane lipids, cholesteryl hemisuccinate (CHS) and phosphatidylcholine (POPC) have unique effects on GPCR structure and function.^17,18^ CHS, a cholesterol analog, and POPC, a common phospholipid, are essential in maintaining membrane organization, fluidity, and stability, contributing to the conformational landscape of membrane proteins.^19,20^ For GPCRs like mGluR2, CHS and POPC are thought to influence receptor activation and stability by modulating the local environment of the transmembrane helices, especially within micelle and bilayer systems.^16,21^ Studies on similar GPCRs, such as mGluR1 and CB1, have demonstrated that cholesterol and CHS stabilize specific receptor conformations, enhance receptor functionality, and maintain the balance between active and inactive states.^22,23^ Cholesterol-rich lipid domains have been shown to act as platforms for mGluR signaling complexes, facilitating signal transduction and receptor localization within cellular membranes.^24^

While structural studies, including X-ray crystallography, have provided valuable static snapshots of GPCRs, complementary techniques like all-atom molecular dynamics (MD) simulations offer dynamic views that reveal conformational flexibility and lipid-mediated modulation at an atomic level.^25–27^ MD simulations enable detailed examination of the conformational transitions that GPCRs undergo in different lipid environments, capturing the influence of lipid composition on structural stability, dynamics, and function. ^28^ Recent MD studies on other GPCRs, such as *β*_2_-adrenergic and A2A receptors, have elucidated critical ligand-binding sites, activation pathways, and conserved interactions that contribute to GPCR stability.^29,30^ However, the molecular mechanisms underlying lipid-mediated modulation of mGluR2, particularly through CHS and POPC in micelle and bilayer systems, remain poorly understood.

In this study, we employed all-atom MD simulations to explore how varying concentrations of CHS (0%, 10%, and 25%) impact mGluR2’s conformational dynamics in both micelle (BLMNG and CHS) and POPC bilayer environments, modeling the receptor in its active and inactive states. Our approach allows for a detailed comparison of mGluR2’s behavior across lipid environments, illuminating the role of CHS in stabilizing the receptor’s transmembrane helices, especially TM1, TM2, and TM3, which constitute the core of the transmembrane domain. We hypothesize that an intermediate concentration of CHS could optimize receptor stability, enhancing the structural rigidity required for effective signaling. Additionally, the comparative analysis of active and inactive states provides insights into state-dependent lipid interactions that may influence mGluR2’s signaling mechanisms.

This work advances our understanding of how lipids modulate G protein-coupled receptor (GPCR) function, offering new perspectives for receptor-specific drug development. Given the physiological importance of mGluR2 within the central nervous system, our results suggest potential avenues for therapeutic intervention in neurological and psychiatric conditions, including schizophrenia, anxiety, and neurodegenerative disorders. By detailing how lipid composition influences the conformational dynamics of mGluR2, our findings highlight the critical role of membrane-protein interactions in preserving GPCR function and lay the groundwork for future strategies in rational drug design.

## Methods

### Simulation Systems

We performed all-atom molecular dynamics (MD) simulations to investigate the conformational dynamics of the metabotropic glutamate receptor 2 (mGluR2) in its active (PDB: 7MTR) and inactive (PDB: 7MTQ) states, embedding each structure in two distinct environments: lipid bilayers and micelles. The lipid bilayer systems consisted of pure 1-palmitoyl-2-oleoyl-sn-glycero-3-phosphocholine (POPC) and heterogeneous bilayers with 10% and 25% cholesteryl hemisuccinate (CHS). For the micelle environment, we used BLMNG (N,N-dimethylglycine betaine-lauryl methane) detergent containing 0%, 10%, or 25% CHS to evaluate CHS-dependent effects on mGluR2 within a micellar system. System setup was carried out using CHARMM-GUI’s Membrane^31–33^ and Micelle Builder modules,^34,35^ which facilitated the placement of mGluR2 into both lipid bilayers and BLMNG-CHS micelles. All systems were solvated with TIP3P water and neutralized with 0.15 M NaCl to mimic physiological conditions. The simulation box dimensions were approximately 187 Å × 187 Å × 187 Å for bilayer systems and 97 Å × 97 Å × 124 Å for micelle systems. Atom counts varied by system type and CHS concentration, with bilayer systems containing between 150,000 and 200,000 atoms and micelle systems comprising approximately 66,000 atoms. The CHARMM36m force field^36,37^ was used for proteins, lipids, and ions, and all simulations were conducted using NAMD 2.14.^38^ Initial energy minimization was performed for 10,000 steps using the conjugate gradient algorithm^39^ to eliminate steric clashes. Equilibration followed CHARMM-GUI protocols under the NVT ensemble, gradually releasing positional restraints over approximately 1 ns. Production simulations were carried out in the NPT ensemble at 310 K, employing a Langevin thermostat (damping coefficient *γ* = 0.5 ps*^−^*^1^) and the Nośe–Hoover Langevin piston method to maintain pressure at 1 atm.^39–41^ Non-bonded interactions were calculated using a cutoff of 10–12 Å, and long-range electrostatics were computed using the particle mesh Ewald (PME) method.^42^ Each simulation was run for 1 *μ*s, resulting in a total simulation time of 36 *μ*s across all systems.

For structural analysis, the transmembrane (TM) helices and loops of mGluR2 were defined as follows: TM1 (residues 565–592), TM2 (597–624), TM3 (629–656), TM4 (676–700), TM5 (724–747), TM6 (758–782), and TM7 (787–810). Intracellular loops (ICLs) and extracellular loops (ECLs) were identified based on previous studies: ICL1 (593–595), ICL2 (657–675), ICL3 (748–757), ECL1 (625–628), ECL2 (701–723), and ECL3 (783–786). To assess conformational stability, root mean square deviation (RMSD) values were calculated for C*α* atoms using VMD’s RMSD Trajectory Tool.^43,44^ Root mean square fluctuation (RMSF) values were also computed to evaluate residue-level flexibility, focusing on TM helices and loop regions. The distance between the centers of mass (COM) of protomer A and protomer B was measured throughout the simulations. Protein structures and trajectories were aligned to the reference structure using C*α* atoms to remove overall translation and rotation. The COM of each protomer was calculated using the measure center function in VMD, and the inter-protomer distance was computed as the magnitude of the vector connecting the two centers. To quantify the relative orientation between protomer A and protomer B, the angle between their principal axes of inertia was calculated for each simulation frame. The backbone atoms of each protomer were selected, and their principal axes were computed using the measure inertia function in VMD. The angle between the second principal axes (representing the longest molecular dimensions) was determined using the dot product followed by the inverse cosine. To ensure consistent interpretation, angles greater than 90*^◦^* were symmetrized using 180*^◦^* − *θ*. RMSD analysis of individual transmembrane helices (TM1–TM7) was performed by aligning each helix to its initial frame using VMD and calculating perframe deviations. Separate selections were made for each helix in both protomers (PROA and PROB), and RMSD values were extracted across all frames to enable comparative structural analysis. Hydrogen bonds were analyzed using the VMD HBond plugin, employing a 3.5 Å distance and 30° angle cutoff. Salt bridges were analyzed using the VMD Timeline plugin with a 4 Å distance cutoff. Salt bridge distances were defined as the shortest distance between donor and acceptor atoms. These simulations provide a comprehensive view of lipid-mediated modulation of mGluR2 in both active and inactive states, highlighting the roles of CHS, POPC, and BLMNG detergent in stabilizing conformational dynamics across micelle and bilayer environments. These findings offer insights relevant to GPCR-targeted drug discovery.

## Results and Discussion

### Structural Stability of mGluR2 Embedded in Micelle Across CHS Concentrations

Our comparative MD simulations revealed notable differences in the structural dynamics of the mGluR2 receptor embedded in micelles between its inactive and active states, as well as across the three studied systems containing 0%, 10%, and 25% CHS. Using RMSD and RMSF analyses, we examined the receptor’s stability and flexibility in both conformational states. In the inactive state, the CHS-free system (0% CHS) displayed the highest RMSD values across the entire protein and in individual protomers A and B, suggesting greater conformational flexibility. In contrast, in the active state, without bound ligand, the system with 25% CHS showed the lowest conformational fluctuations.

RMSD values for the transmembrane (TM) domain remained below 6 Å during the inactive state simulations in systems with 10% and 25% CHS, as well as in the active state simulations with 0% and 10% CHS. This indicates minimal deviation from the initial antagonist-bound crystal structure and suggests structural stability after agonist dissociation in the active state (Figure 1 and S1-S3). These consistent RMSD values reflect sustained rigidity, supporting the notion that CHS at these concentrations helps maintain stable conformations of the TM domain, regardless of the receptor’s activation state.

**Fig. 1.**
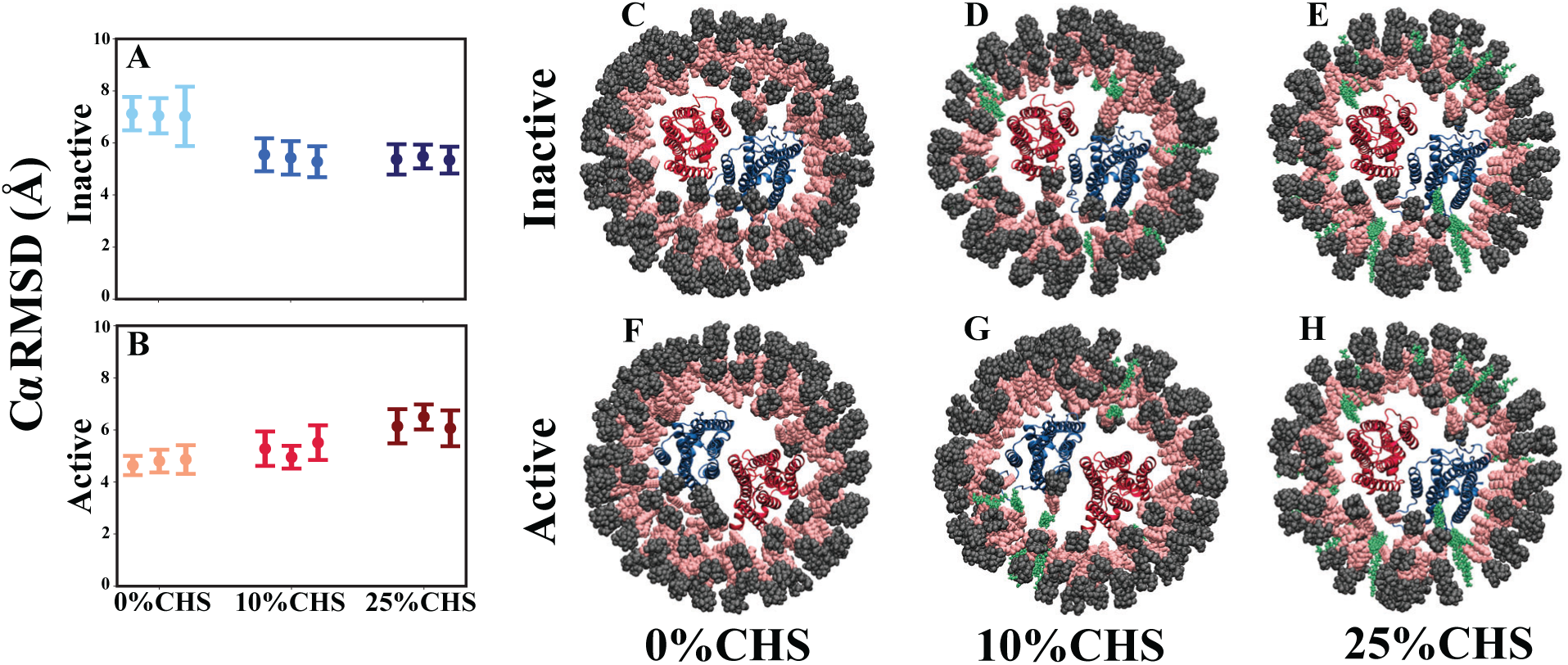
Structural stability of mGluR2 embedded in micelle in active and inactive states. (A) Average C*α* RMSD of mGluR2 in the inactive state across CHS concentrations (0%, 10%, 25%) from three replicates. (B) Average C*α* RMSD of mGluR2 in the active state under the same conditions. (C–E) Representative snapshots of mGluR2 in the inactive state at 0%, 10%, and 25% CHS, respectively. (F–H) Snapshots in the active state at 0%, 10%, and 25% CHS, respectively. CHS molecules are shown in green, with the bilayer and receptor colored distinctly. Error bars in (A) and (B) represent standard deviations, reflecting CHS effects on receptor stability.

Conversely, the elevated RMSD in the CHS-free inactive system indicates greater flexibility and conformational diversity, possibly reflecting a broader ensemble of intermediate structures relevant to state transitions in ligand-free conditions. Meanwhile, the reduced RMSD in the active state with 25% CHS supports the hypothesis that higher CHS levels constrain the receptor in an active-like conformation, potentially facilitating downstream processes such as G protein coupling. This structural rigidity implies that CHS reinforces the active state and reduces the receptor’s tendency to adopt alternative conformations.

Per-residue RMSF profiles revealed state-dependent and CHS concentration-dependent differences in receptor flexibility (Figure 2). Flexibility was generally elevated in extracellular and intracellular loops compared to the more rigid TM cores. However, the degree of flexibility varied with CHS concentration, receptor state, and protomer identity.

**Fig. 2.**
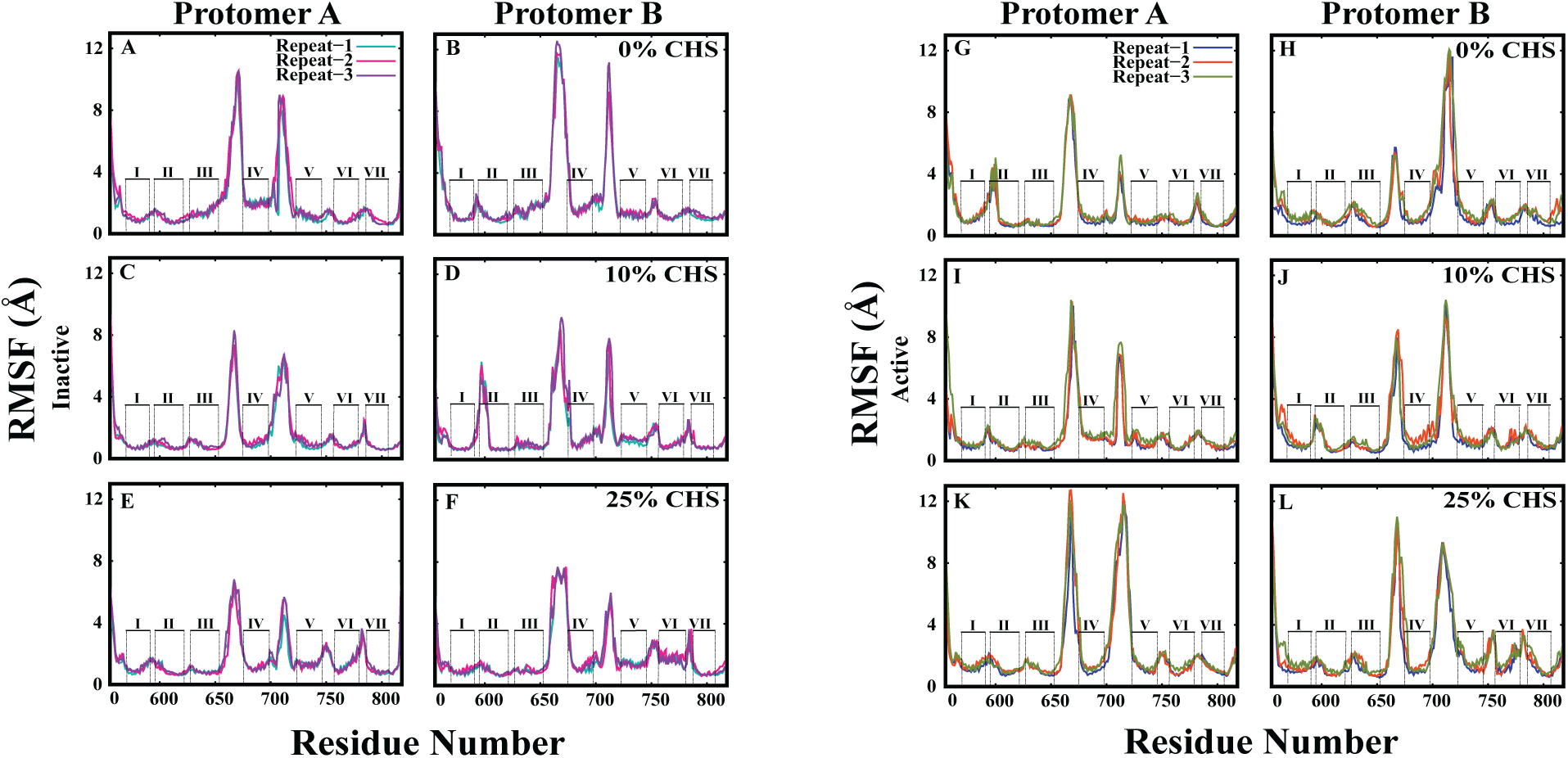
Root Mean Square Fluctuation (RMSF) analysis of mGluR2 in a micelle across CHS concentrations. Left panels: RMSF profiles for Protomers A and B in the inactive state; right panels: active state. Each panel shows residue-wise fluctuations at 0%, 10%, and 25% CHS across three independent replicates. Transmembrane helices (I–VII) are indicated. Notable differences occur in TM helices IV and VI, where CHS modulates flexibility in a state- and protomer-dependent manner.

In the inactive state, the TM3-4 and TM4-5 interhelical loops were most flexible in the CHS-free system, followed by 10% CHS, and least flexible at 25% CHS. Interestingly, this trend was reversed in the active state, with increased loop flexibility at higher CHS levels.

A detailed comparison across interhelical loops revealed specific differences between protomers and states. For instance, in the inactive state, the loop between TM1 and TM2 in protomer B exhibited peak flexibility at 10% CHS. In the active state, this same loop in protomer A was most flexible without CHS and showed reduced flexibility with increasing CHS concentration. The TM3-4 loop followed a similar trend: most flexible in the CHS-free inactive state and less so with more CHS, but in the active state, the trend was reversed. For TM4-5, the same pattern held in protomer A, while in protomer B, it remained consistent across states. TM6-7 loop flexibility increased with CHS in the inactive state but reversed in protomer A during the active state, whereas protomer B retained the initial trend.

The pronounced flexibility in the CHS-free inactive system suggests an increased capacity of mGluR2 to explore alternative conformations, possibly a functional advantage for GPCRs that transition between states. As a class C GPCR, mGluR2 relies on such flexibility for ligand recognition and signal transduction. CHS appears to modulate the receptor’s energy landscape: at low concentrations, it permits dynamic transitions, whereas at higher concentrations, it enforces rigidity, particularly in the active state, potentially stabilizing functionally relevant conformations.

Additionally, differences between protomers A and B highlight asymmetric CHS effects, which may influence cooperative behavior in the receptor dimer. The opposite trends in RMSF between active and inactive states, especially in key interhelical loops, suggest CHS modulates not just static stability but also dynamic transitions. These results emphasize cholesterol’s nuanced role in shaping membrane protein behavior, reinforcing the concept that lipid environment critically determines GPCR conformational landscapes and signaling capacities.

### Cholesteryl-Mediated Fine-Tuning of mGluR2 Dimer Stability and Signaling

The effects of CHS concentration on inter-protomer distance and orientation in mGluR2 reveal a state-dependent structural modulation that may influence receptor function. In the inactive state, inter-protomer distance increased with CHS concentration, with the CHS-free system exhibiting the shortest distance, followed by 10% CHS, and the largest distance observed at 25% CHS (Figure 3 and S4). This suggests that higher CHS concentrations promote a more expanded dimeric conformation, potentially affecting the receptor’s ability to transition into an active state. In contrast, in the active state, this trend was reversed, with the shortest inter-protomer distance observed in the CHS-free system and similar, slightly larger distances at 10% and 25% CHS. This compaction of the dimer interface in the active state at lower CHS levels may reflect a cholesterol-dependent stabilization of the active-like conformation, potentially influencing downstream signaling efficiency.

**Fig. 3.**
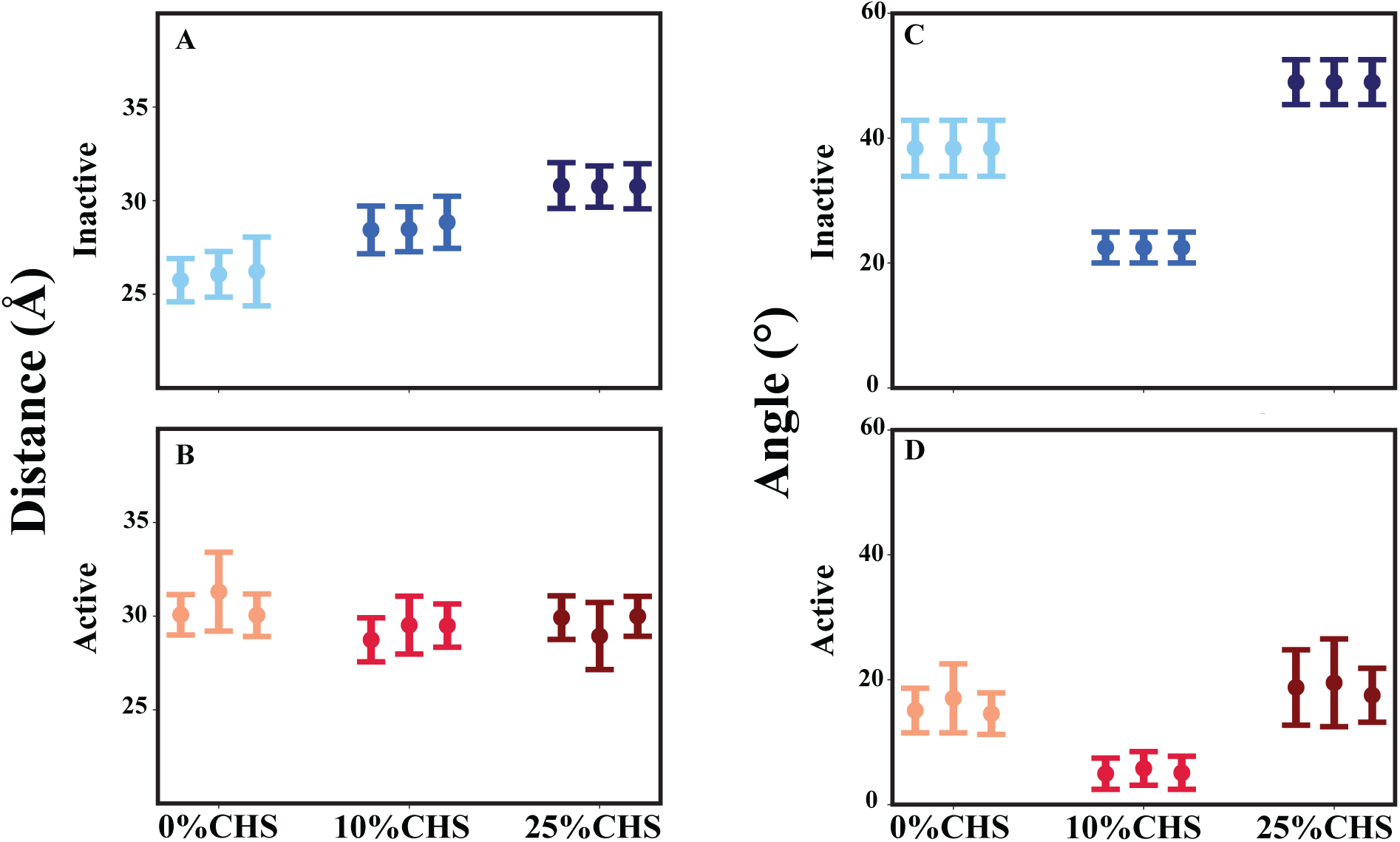
Interprotomer Distance and Angle Analysis of mGluR2 in Active and Inactive States. (A) Average interprotomer distance between Protomer A and Protomer B of the mGluR2 receptor in the inactive state under varying CHS concentrations (0%, 10%, and 25%), calculated from three independent simulations. (B) Average interprotomer distance for the active state under the same CHS conditions. (C) Average interprotomer angle between Protomer A and Protomer B in the inactive state, highlighting structural differences under varying CHS concentrations. (D) Average interprotomer angle for the active state. The error bars represent standard deviations, illustrating the impact of CHS concentrations on the structural stability and relative orientation of the two protomers in micelle environments.

The inter-protomer angle also exhibited a nonlinear dependence on CHS concentration, indicating a cholesterol-driven reorganization of the dimeric structure. In the inactive state, the 10% CHS system showed the lowest inter-protomer angle, while the CHS-free and 25% CHS systems displayed larger angles. A similar trend was observed in the active state, with the 10% CHS system maintaining the lowest angle, while both the 0% and 25% CHS systems exhibited higher values (Figure 3 and S5). These observations suggest that at an intermediate CHS concentration, mGluR2 adopts a conformation that balances structural stability and flexibility, potentially optimizing ligand-induced activation and G-protein coupling.

Functionally, these results align with mGluR2’s role as an obligate dimer, where cholesterol-dependent changes in inter-protomer distance and orientation can modulate signal transduction efficiency. The increased inter-protomer distance at higher CHS concentrations in the inactive state, coupled with reduced loop flexibility (as seen in RMSF analyses), suggests that CHS may stabilize an inactive-like conformation, limiting the receptor’s capacity to transition to an active state. In the active state, CHS appears to restrict excessive fluctuations, potentially maintaining a structurally stable active conformation that could support interactions with intracellular signaling partners such as G-proteins.

This cholesterol-induced fine-tuning of receptor architecture may be essential in regulating mGluR2’s functional dynamics, influencing its ability to adopt conformations that favor activation or reinforce the inactive state depending on the cellular environment. These findings indicate that CHS plays a dual role in mGluR2 function: it constrains inter-protomer movement to stabilize the active state and allows greater conformational diversity in the inactive state, possibly to facilitate ligand binding and state transitions. This cholesterol-dependent modulation underscores the importance of membrane composition in GPCR signaling and suggests that CHS levels could regulate mGluR2 activity in varying physiological or pathological conditions.

### State-Dependent Structural Rearrangements of Individual TM Helices in mGluR2

To shed more light on the state-dependent conformational changes in mGluR2, we conducted a detailed analysis of RMSD variations across individual transmembrane (TM) helices in both the inactive and active states. These findings provide insights into the differential stability and flexibility of specific helices, revealing CHS-mediated structural effects that may influence receptor function.

In the inactive state, TM1 remained structurally stable across all CHS concentrations in both protomers A and B, suggesting minimal conformational fluctuations. Similarly, TM5 and TM7 displayed consistent stability across all systems, indicating their role as anchoring elements that are not significantly affected by CHS in the inactive conformation. TM2 in protomer A remained stable under all conditions, whereas in protomer B, 10% CHS induced the highest flexibility, implying localized destabilization effects. TM3 and TM4 exhibited the greatest flexibility in the absence of CHS, with 0% CHS showing the highest RMSD values (Figure 4 and and S6). This suggests that CHS-free conditions allow these helices to explore a broader conformational space, potentially affecting receptor activation, which aligns with findings from other GPCR studies indicating that TM3 undergoes substantial structural rearrangements upon activation.^45–47^

**Fig. 4.**
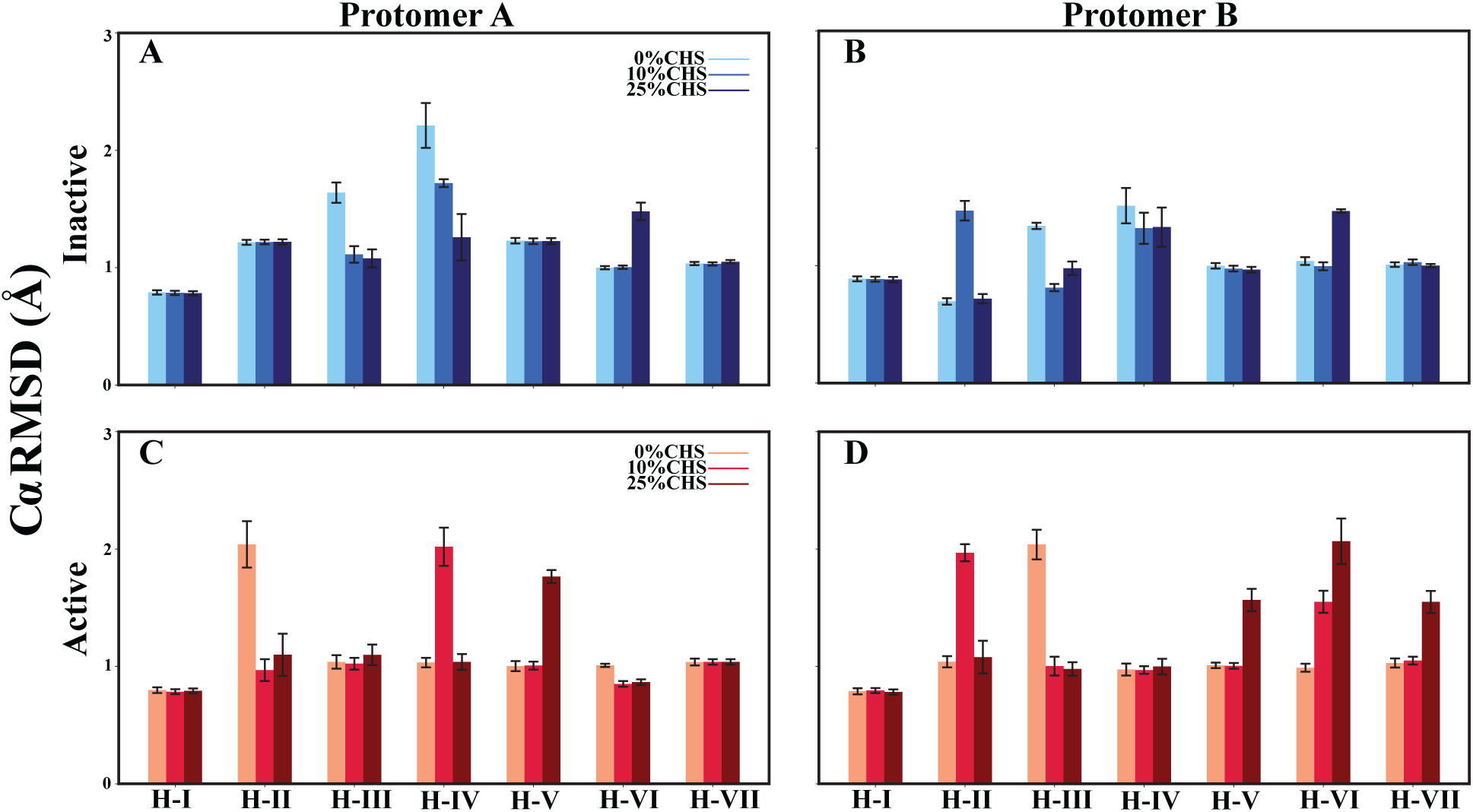
Mean and Standard Deviation of C*α* RMSD for Transmembrane Helices (H-I to H-VII) in mGluR2 Under Varying CHS Concentrations. C*α* RMSD values are shown for individual helices of Protomer A (panels A, C) and Protomer B (panels B, D) of mGluR2 in both the inactive (A–B) and active (C–D) states, embedded in a micelle environment. Bars represent the mean RMSD across three independent replicas, with error bars denoting standard deviation. Three CHS concentrations (0%, 10%, and 25%) were tested to evaluate the effect of cholesterol on the structural flexibility of each transmembrane helix. The results indicate CHS-dependent changes in helix-specific stability, particularly in the active state, where selected helices exhibit enhanced fluctuations at higher CHS concentrations.

A study on mGluR2 using FRET sensors revealed that activation involves conformational changes correlated with the dynamics of TM3, demonstrating its crucial role in allosteric modulation and receptor activation.^48^ Other investigations into GPCRs similarly highlight the involvement of TM3 in activation processes.^49–52^ Moreover, CHS has been shown to stabilize GPCRs by forming bicelle-like micelle architectures with detergents, as observed in the NOP receptor.^53^ smFRET studies further indicate that cholesterol shifts the equilibrium between active and inactive states of mGluR2, underscoring its role in fine-tuning receptor function.^54^

Notably, TM6 exhibited the highest RMSD at 25% CHS in both protomers, suggesting that elevated CHS concentrations induce structural rearrangements that may stabilize specific receptor states. These observations align with previous findings demonstrating that lipid interactions with TM6 can influence GPCR activation pathways by modulating helical flexibility and conformational transitions.^14,26,55^

In the active state, TM1 again remained highly stable across all CHS conditions in both protomers. TM2 exhibited differential stability patterns, with protomer A showing the highest RMSD at 0% CHS, while protomer B displayed the greatest flexibility at 10% CHS. TM3 exhibited minimal fluctuations in protomer A, whereas in protomer B, the CHS-free system showed the highest RMSD, indicating that CHS modulates the conformational landscape differently across protomers (Figure 4 and and S7). This supports findings that TM3 stability is key to receptor activation. ^56–58^

TM4 demonstrated state-dependent fluctuations: protomer A had the highest RMSD at 10% CHS, while protomer B showed similar stability across all CHS levels. TM5 displayed the highest RMSD at 25% CHS in both protomers, suggesting that elevated CHS concentrations introduce increased flexibility in this helix. TM6 followed a similar pattern, with 0% CHS yielding the highest RMSD in protomer A, and 25% CHS causing the greatest fluctuations in protomer B. TM7 showed asymmetric behavior: while remaining stable in protomer A, it displayed the highest RMSD at 25% CHS in protomer B. This may reflect a CHS-mediated modulation of TM7 mobility, consistent with Family A GPCRs where TM7 rearrangements are essential for activation.^17,23,47^

The interplay between TM1, TM2, TM6, TM7, TM3, TM5, and the loop between TM4 and TM5 plays a pivotal role in shaping mGluR2’s structural transitions. These helices undergo CHS-dependent rearrangements that influence overall receptor stability and interhelical communication. Cholesterol binds at specific high-affinity sites—particularly around TM5-TM7—stabilizing the receptor and enhancing activation efficiency. These binding sites are evolutionarily conserved, emphasizing their critical role in allosteric regulation.^59^

Further analysis reveals distinct CHS-dependent behaviors: in the inactive state, TM6 exhibits the highest RMSD at 25% CHS, suggesting CHS-induced destabilization in the resting conformation. In contrast, in the active state, TM6 responds asymmetrically—0% CHS causes the highest RMSD in protomer A, while 25% CHS increases fluctuations in protomer B. This may reflect cholesterol’s dual role in stabilizing one protomer while promoting flexibility in the other, potentially fine-tuning activation.^16,59^

TM5 remains stable in the inactive state across all CHS levels, but shows highest flexibility at 25% CHS in the active state. TM4, particularly in protomer A, is destabilized at 10% CHS. TM3 again shows increased fluctuations in CHS-free conditions, suggesting that CHS helps constrain this helix into stable conformations across both states. TM2 follows a state- and protomer-specific pattern: 0% CHS induces high RMSD in protomer A (active state), while 10% CHS does so in protomer B. Beyond RMSD trends, CHS-dependent hydrogen bonds involving TM1 with TM2 and TM6 with TM7 contribute to structural stabilization. Certain CHS concentrations enhance inter-helical coupling, while others lead to bond disruption, potentially fine-tuning receptor activation. Additionally, the interaction between TM3 and the TM4/5 loop provides a structural link between cholesterol binding and receptor flexibility, suggesting a mechanistic pathway through which CHS regulates activation transitions.

### Cholesterol-Dependent Modulation of Hydrogen Bonding in mGluR2

To further elucidate the structural and functional role of cholesterol (CHS) in mGluR2 allosteric regulation, we analyzed the formation and stability of interhelical hydrogen bonds in both inactive and active states. Cholesterol interacts with specific binding sites within GPCRs, modulating receptor conformational flexibility and transitions between states. Hydrogen bonds between transmembrane (TM) helices play a crucial role in maintaining receptor integrity, stabilizing specific conformations, and facilitating these state transitions. Our findings indicate that CHS concentration modulates key interhelical hydrogen bond interactions in a state- and protomer-dependent manner, potentially through direct binding at cholesterol interaction sites, influencing receptor dynamics and activation efficiency.

The TM1–TM2 interaction, which is critical for receptor topology, exhibited state-dependent modulation through the ASN593–ARG603 hydrogen bond. In the inactive state, this bond remained consistently formed in Protomer A across all CHS conditions (0%, 10%, and 25%), suggesting that this interaction is structurally stable and largely independent of CHS influence. However, in Protomer B, the bond was present at 0% and 25% CHS but absent at 10% CHS, indicating that an intermediate CHS concentration disrupts this interaction, potentially due to CHS-mediated perturbations at binding sites near TM1 and TM2. In the active state, the TM1–TM2 bond became more dynamic, with CHS influencing its formation and disruption. In Protomer A, the bond was absent at 0% CHS, formed at 10% CHS, and lost again at 25% CHS, whereas in Protomer B, the bond was present at 0% and 25% CHS but absent at 10% CHS (Figure 5), suggesting that CHS fine-tunes receptor activation by selectively stabilizing or destabilizing TM1–TM2 interactions.

**Fig. 5.**
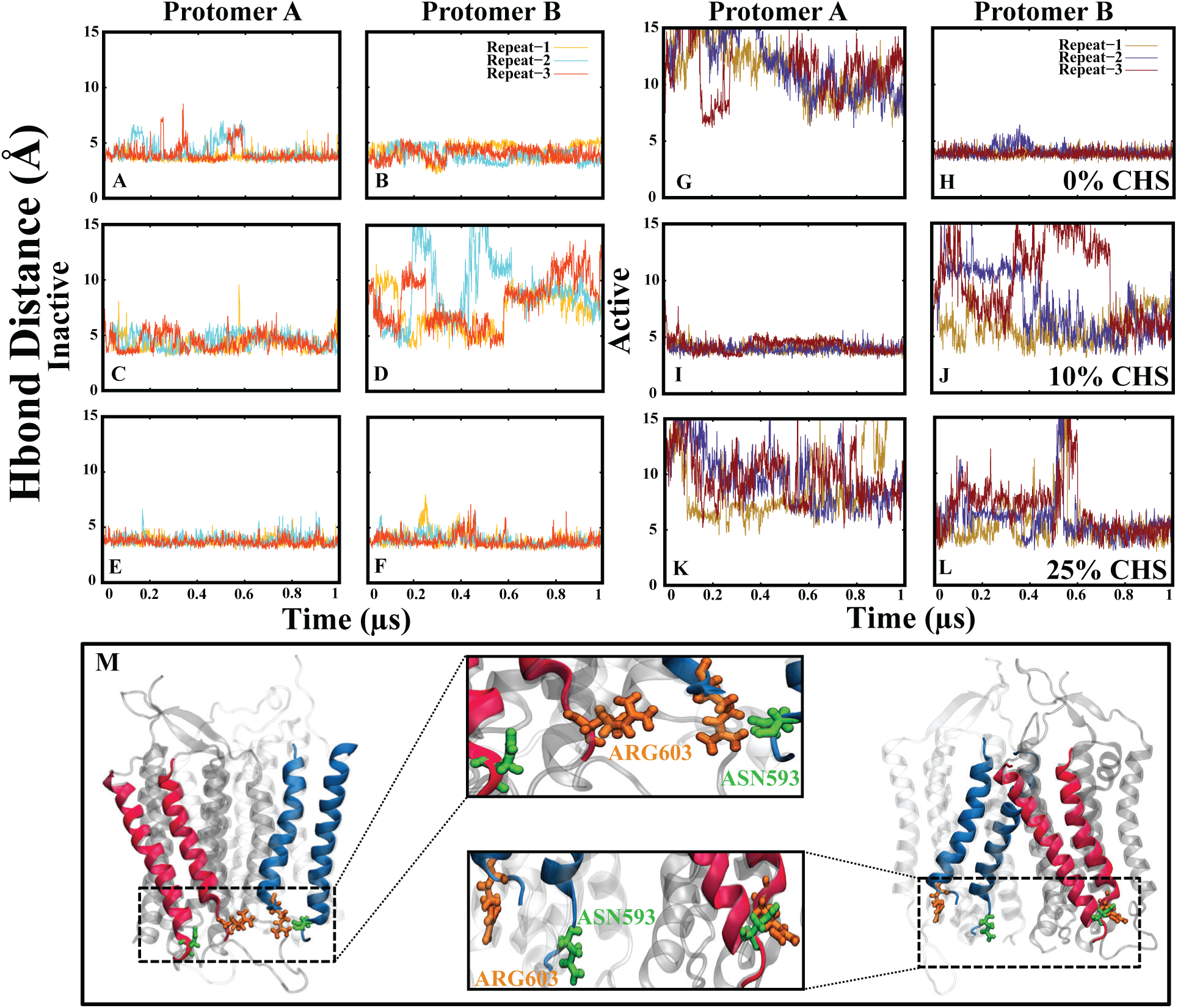
Hydrogen bond interaction between ARG603 and ASN593 stabilizes the TM1 and TM2 helices in the inactive and active states of mGluR2 embedded in a bilayer. Panels (A, B) show the time series of the hydrogen bond distances between the C*α* atoms of ARG603 (TM2) and ASN593 (TM1) at 0% CHS in Protomer A and Protomer B in the inactive state. Panels (C, D) and (E, F) display the corresponding distances at 10% CHS and 25% CHS in Protomer A and Protomer B in the inactive state, respectively. Panels (G, H) illustrate the hydrogen bond distances at 0% CHS in Protomer A and Protomer B in the active state, while panels (I, J) and (K, L) depict the data at 10% CHS and 25% CHS in Protomer A and Protomer B in the active state, respectively. Panel (M) presents a structural representation of the ARG603–ASN593 hydrogen bond interactions within the TM1–TM2 interface, highlighting their spatial positioning within the receptor’s transmembrane helices.

The TM3–loop (TM4–TM5) interaction, mediated by the THR629–THR705 hydrogen bond, showed a CHS-dependent stabilization trend. In the inactive state, this bond was absent at 0% CHS in both protomers, indicating a high degree of TM3 flexibility in cholesterol-free environments. At 10% CHS, the bond formed rapidly in Protomer B but showed a delayed formation in Protomer A, highlighting differential cholesterol stabilization effects. At 25% CHS, this bond remained consistently present in both protomers (Figure 6), reinforcing the idea that higher cholesterol concentrations induce structural rigidity, potentially limiting TM3 movement by stabilizing interactions at CHS binding hotspots. In the active state, the TM3–loop bond displayed a similar protomer-asymmetric response. At 0% CHS, the bond remained absent in both protomers. At 10% CHS, the bond failed to form in Protomer A but appeared after 0.5 ns in Protomer B. A similar trend was observed at 25% CHS, where the bond remained absent in Protomer A but was stable in Protomer B.

**Fig. 6.**
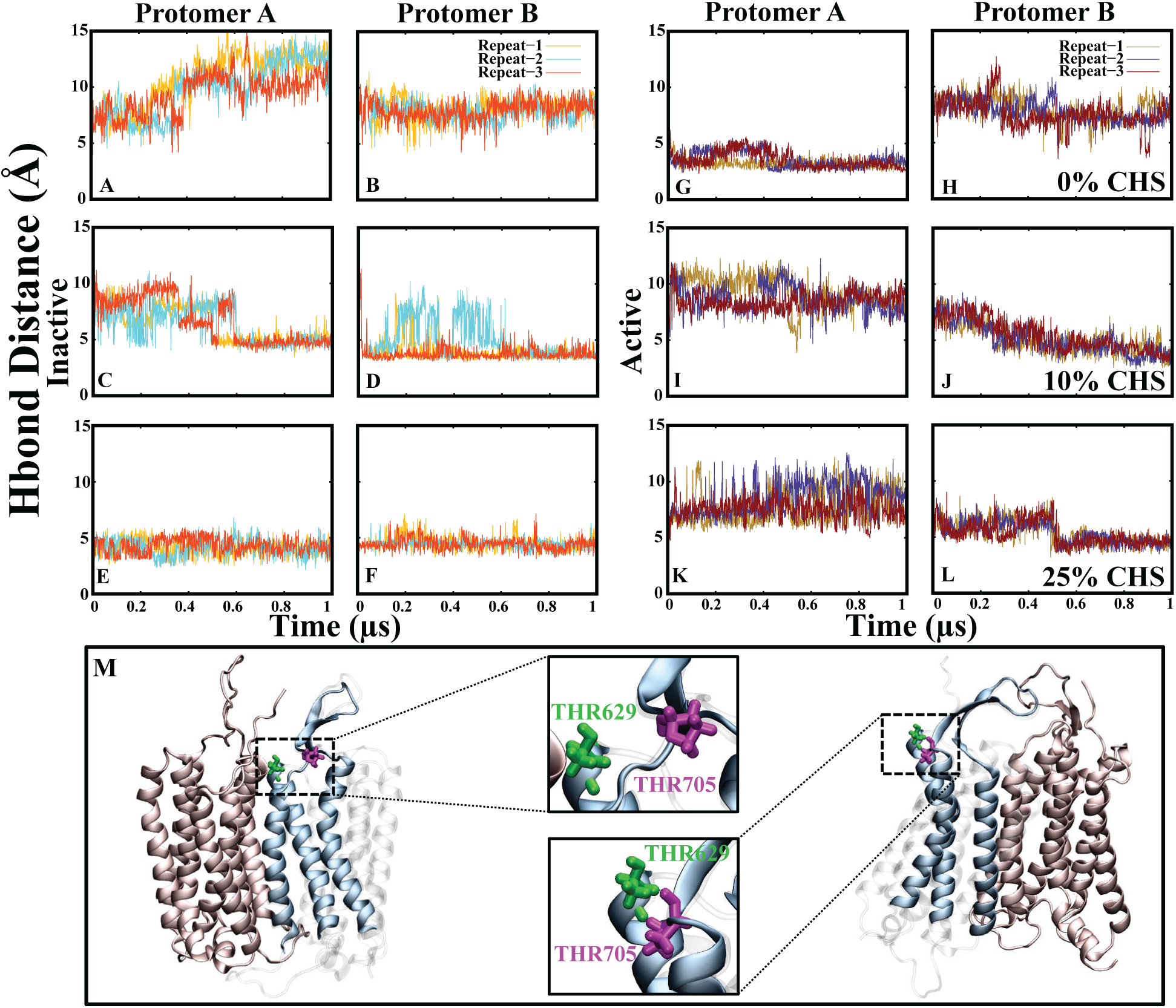
Hydrogen bond interaction between THR629 and PRO703 stabilizes the TM3–loop (TM4–TM5) region in the inactive and active states of mGluR2 embedded in a bilayer. Panels (A, B) represent the time series of the hydrogen bond distances between the C*α* atoms of THR629 (TM3) and THR705 (loop between TM4–TM5) at 0% CHS in Protomer A and B in the inactive state. Panels (C, D) and (E, F) show the corresponding distances at 10% CHS and 25% CHS. Panels (G, H) illustrate the hydrogen bond distances at 0% CHS in the active state. Panels (I, J) and (K, L) depict the data at 10% and 25% CHS in the active state. Panel (M) provides a structural representation of the THR629–THR705 hydrogen bond.

The TM6–TM7 interaction, which plays a key role in stabilizing the receptor’s transmembrane core, exhibited a CHS-sensitive formation pattern through the TRP773–SER801 hydrogen bond. At 0% CHS, this bond was stable in both protomers, reinforcing the idea that cholesterol-free environments promote tighter TM6–TM7 interactions. At 10% CHS, the bond formed after a delay of 0.2 *μ*s, suggesting that CHS binding at nearby sites modulates receptor flexibility by stabilizing key transmembrane contacts. At 25% CHS, the bond was completely absent in both protomers, indicating that excess CHS disrupts TM6–TM7 interactions, potentially through excessive stabilization at allosteric binding sites.^59^ In the active state, CHS exerted asymmetric control over TM6–TM7 interactions. In Protomer A, the bond did not form at 0% CHS, while in Protomer B, it remained intact. At 10% CHS, the bond formed later in the simulation in Protomer A but appeared earlier in Protomer B (Figure 7).

**Fig. 7.**
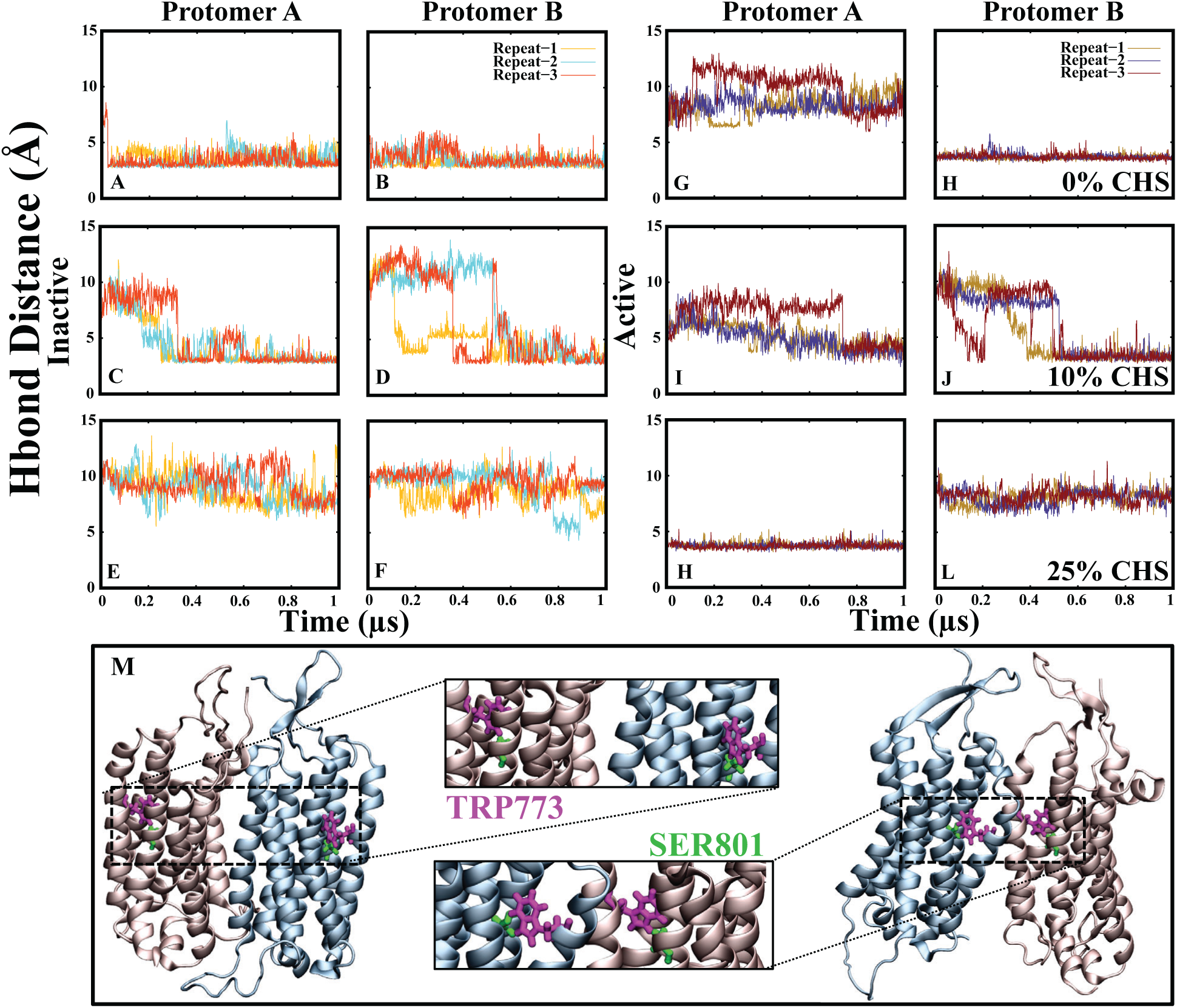
Hydrogen bond interaction between TRP773 and SER801 stabilizes the TM6 and TM7 helices in the inactive and active states of mGluR2 embedded in a bilayer. Panels (A, B) represent the time series of the hydrogen bond distances between the C*α* atoms of TRP773 (TM6) and SER801 (TM7) at 0% CHS in Protomer A and Protomer B in the inactive state. Panels (C, D) and (E, F) show the corresponding distances at 10% CHS and 25% CHS. Panels (G, H) illustrate the hydrogen bond distances at 0% CHS in the active state. Panels (I, J) and (K, L) depict the data at 10% and 25% CHS in the active state. Panel (M) provides a structural representation of the TRP773–SER801 hydrogen bond within the TM6–TM7 interface.

### Salt Bridge Dynamics and Their Relationship to Protomer Distance and Angle in mGluR2

Our simulations revealed a dynamic network of salt bridge interactions in mGluR2, exhibiting distinct state-dependent formation and stability across different cholesteryl (CHS) conditions (Figure 8). These salt bridges play a crucial role in stabilizing receptor conformations, influencing inter-protomer distance and angle fluctuations, and ultimately shaping the receptor’s conformational landscape. Electrostatic interactions, particularly salt bridges, have been recognized as critical determinants of GPCR state transitions, acting as molecular switches that influence receptor stability and activation. In mGluR2, our findings suggest that the formation and disruption of these interactions are closely linked to CHS concentration, modulating receptor compaction and flexibility in a state-dependent manner.

**Fig. 8.**
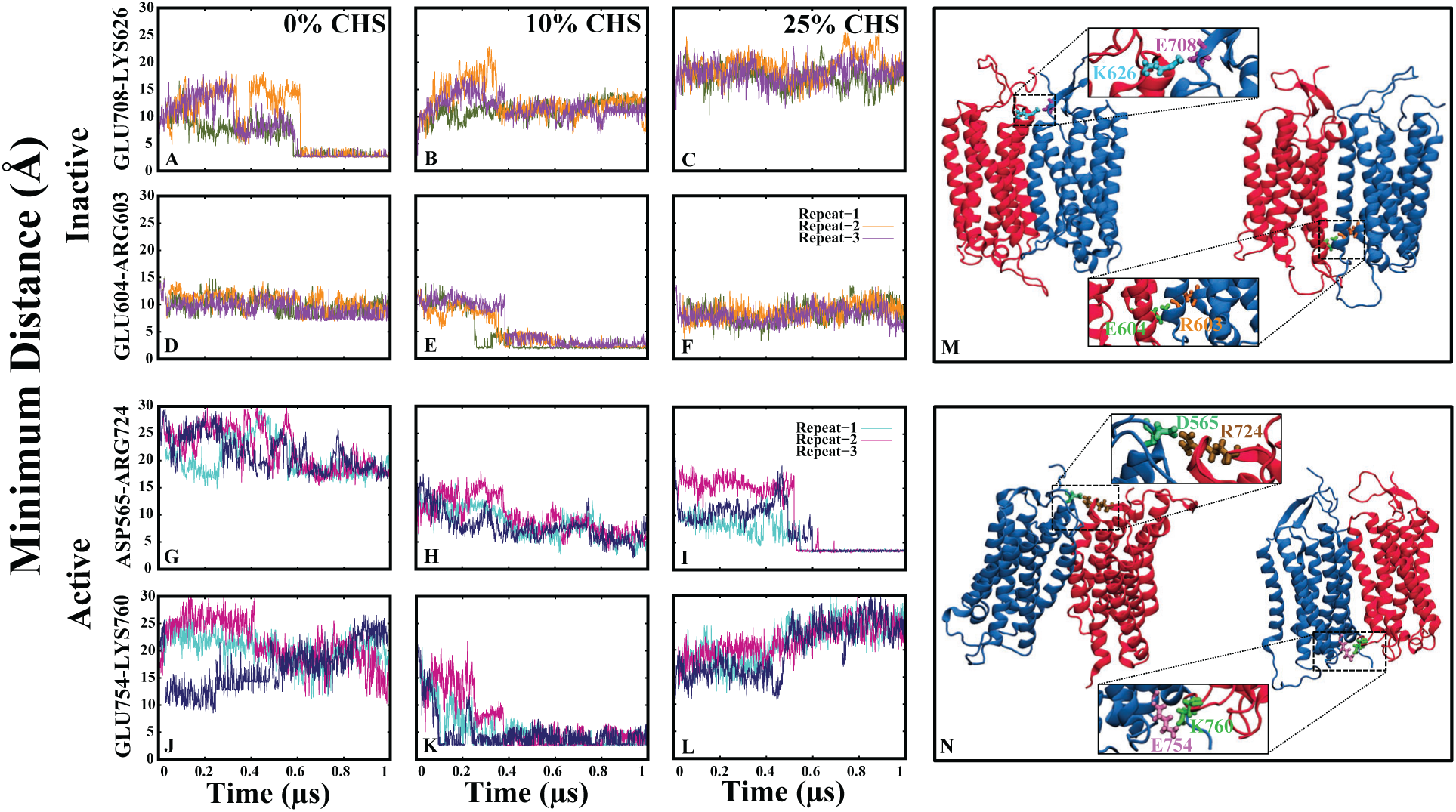
Minimum distance analysis of salt-bridge interactions in mGluR2 embedded in a micelle. Panels (A, B, C) show the time series of the minimum distances between Glu708–Lys626 in the inactive state at 0%, 10%, and 25% CHS, respectively. Panels (D, E, F) display the minimum distances of the Glu604–Arg603 interaction in the inactive state at 0%, 10%, and 25% CHS. Panels (G, H, I) represent the minimum distances of the Asp565–Arg724 interaction in the active state at 0%, 10%, and 25% CHS. Panels (J, K, L) show the minimum distance analysis of the Glu754–Lys760 interaction in the active state at 0%, 10%, and 25% CHS. Panels (M, N) provide structural representations of the salt-bridge interactions, highlighting key residue pairs within the micelle environment. The three colored lines in each panel correspond to independent simulation replicates.

In the inactive state, we identified a salt bridge between Glu708 (loop between Helix 4 and Helix 5 on Protomer B) and Lys626 (loop between Helix 2 and Helix 3 on Protomer A) that formed at the end of the simulation in the CHS-free condition (0% CHS). The formation of this interaction coincides with a reduction in inter-protomer distance, suggesting that electrostatic forces contribute to maintaining a more compact receptor conformation in the absence of CHS. The stabilization of this interaction late in the simulation indicates that under CHS-free conditions, the receptor may rely on these electrostatic contacts to preserve its structural integrity. This aligns with previous computational studies showing that Asp/Glu···Arg salt bridges are generally more stable than Asp/Glu···Lys interactions, even in environments with moderate polarity, reinforcing the idea that CHS-free conditions may promote salt bridge formation through altered dielectric properties.^60^

Another significant inactive-state salt bridge emerged at 10% CHS, where Glu604 (Helix 2, protomer A) and Arg603 (Helix 2, protomer B) formed a stable interaction after 0.3 ns. This salt bridge aligned with a decrease in the inter-protomer angle, suggesting that CHS modulates receptor compaction via inter-protomer electrostatic interactions. Interestingly, Arg603 in protomer B did not form a hydrogen bond with Phe589 at 10% CHS; instead, it formed a salt bridge with Glu604 in protomer A. This finding indicates that CHS alters interprotomer electrostatic interactions in a concentration-dependent manner, which is consistent with previous studies on eIF2*α* kinases, where functionally important salt bridge interactions regulate stability and activity, as well as membrane protein oligomerization studies, where specific salt bridges drive protomer assembly and stabilization.^61,62^ Additionally, Helix 2, which exhibited the highest movement at the beginning of the simulation, showed a gradual reduction in flexibility upon salt bridge formation in protomer B, indicating a potential stabilizing role of this electrostatic contact in restraining domain motions.

In the active state, a distinct set of salt bridge interactions emerged, further linking electrostatic rearrangements to receptor activation. At 25% CHS, Asp565 (Helix 1, Protomer A) formed a stable salt bridge with Arg724 (Helix 5, Protomer B) around the midpoint of the simulation and remained intact throughout. The formation of this interaction coincided with reduced movement in Helix 5, suggesting that salt bridge formation at 25% CHS stabilizes the receptor at later stages rather than during early activation dynamics. Moreover, these fluctuations correlate with a higher inter-protomer distance at 0% CHS, reinforcing the notion that CHS concentration differentially regulates protomer alignment and receptor stability. These findings align with previous studies showing that salt bridges play a crucial role in GPCR activation and structural stability, as demonstrated in the gonadotropin-releasing hormone receptor (GnRHR), where a salt bridge between transmembrane segments 2 and 3 was essential for receptor activation and ligand specificity.^63^ Additionally, research on GPCRs in nanodiscs and LMNG/CHS micelles has highlighted the role of lipid environments in stabilizing key receptor interactions, particularly at lipid–water interfaces.^64^

Finally, in the active state at 10% CHS, we observed the formation of a salt bridge between Glu754 (loop between Helix 5 and Helix 6, Protomer A) and Lys760 (Helix 6, Protomer B). This salt bridge aligns with a decrease in the inter-protomer angle at 10% CHS, suggesting that electrostatic stabilization between Helices 5 and 6 contributes to structural compaction at intermediate CHS concentrations. Given that salt bridge formation coincided with changes in the inter-protomer angle, it is likely that CHS modulates receptor flexibility by promoting electrostatic contacts that stabilize the activated receptor conformation. This behavior is consistent with previous findings in rhodopsin activation, where interhelical salt bridge rearrangements are essential for receptor conformational transitions and signal propagation, further emphasizing the role of electrostatic interactions in GPCR activation. ^65^ These findings highlight a state- and CHS-dependent salt bridge network that influences inter-protomer distance, angle fluctuations, and domain stability in mGluR2. Low CHS (0%) favors compact inactive-state conformations through intra-loop salt bridge formation, while moderate CHS (10%) promotes targeted stabilization between Helices 2, 5, and 6. Higher CHS (25%) contributes to activation stabilization but allows flexibility for later transitions. These trends align with prior observations in GPCRs, where salt bridge rearrangements are recognized as key determinants of receptor activation and state transitions.^65,66^ The correlation between salt bridge formation and distance/angle metrics in mGluR2 underscores the functional importance of electrostatic interactions in regulating receptor conformational landscapes.

### Structural Stability of mGluR2 Embedded in a Bilayer Across CHS Concentrations

To expand upon our previous analysis of mGluR2 embedded in micelles, we examined how embedding the receptor in a lipid bilayer affects its conformational dynamics across inactive and active states at 0%, 10%, and 25% CHS. Using RMSD and RMSF analyses, we evaluated structural flexibility and stability under each condition.

In the inactive state, the CHS-free system (0%) exhibited the highest RMSD values, indicating greater flexibility in the absence of CHS. Conversely, in the active state, the system with 10% CHS displayed the highest conformational fluctuations, suggesting that intermediate CHS levels induce more pronounced structural rearrangements than either 0% or 25% CHS.

While 10% and 25% CHS stabilized the inactive conformation, the enhanced dynamics observed in the active state at 10% CHS (Figure 9 and S8-S10) imply that this concentration may enhance receptor flexibility rather than stabilizing a specific active-like structure. This observation is consistent with prior computational studies on the *β*-adrenergic receptor and serotonin1A receptor, where intermediate cholesterol or CHS levels increased receptor flexibility.^17,67^ At 25% CHS, the active state adopted a more constrained structure, suggesting that higher CHS levels promote stabilization, as similarly observed in studies of rhodopsin and CX3CR1.^68^ These findings support the idea that 10% CHS represents a threshold concentration that perturbs, rather than reinforces, receptor stability—possibly facilitating activation or intermediate conformational sampling. Similar mechanisms have been reported in CCR3 and cannabinoid receptors, where intermediate lipid concentrations increase flexibility and promote activation.^69,70^

**Fig. 9.**
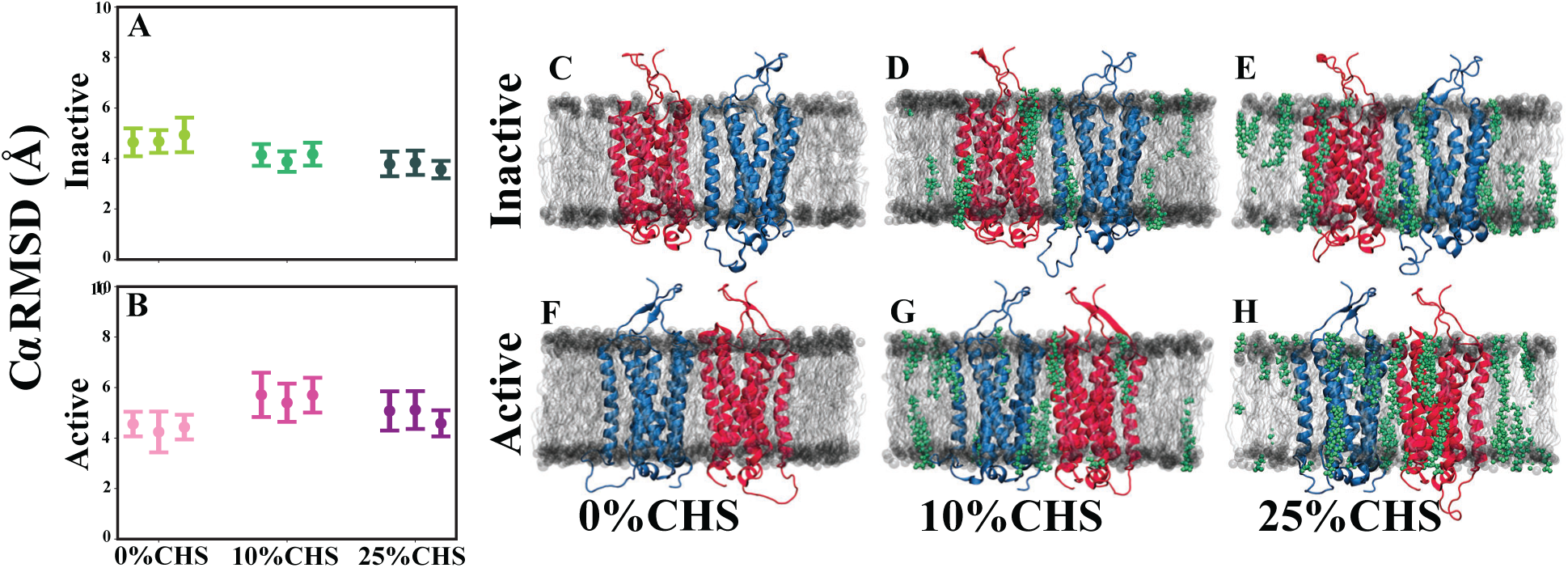
Structural stability of the mGluR2 receptor in active and inactive states. (A) Average C*α* RMSD of mGluR2 in the inactive state under varying CHS concentrations (0%, 10%, and 25%), calculated from three independent simulations. (B) Average C*α* RMSD of mGluR2 in the active state under the same CHS conditions. (C–E) Representative snapshots of mGluR2 embedded in a bilayer in the inactive state at 0%, 10%, and 25% CHS, respectively. (F–H) Representative snapshots of mGluR2 in the active state at 0%, 10%, and 25% CHS, respectively. CHS molecules are shown in green, while the bilayer environment and receptor structure are visualized in distinct colors. The error bars in (A) and (B) represent standard deviations, highlighting the effect of CHS concentrations on receptor stability within bilayer environments.

The RMSD of the TM domain remained below 6 Å throughout the inactive state at 10% and 25% CHS, as well as in the active state for both 0% and 25% CHS. However, the elevated RMSD at 10% CHS in the active state suggests a more dynamic conformational ensemble. This supports a dual model where moderate CHS concentrations facilitate conformational transitions and higher CHS levels enforce a more stable active-like state. These data reinforce the concept that CHS modulates the receptor energy landscape in a concentration-dependent manner, potentially influencing downstream signaling via structural stabilization or flexibility.

To further investigate the molecular basis for these RMSD trends, we analyzed per-residue RMSF profiles (Figure 10). Loop regions consistently exhibited higher fluctuations than TM helices, but the bilayer environment revealed CHS-dependent effects distinct from the micelle system.

**Fig. 10.**
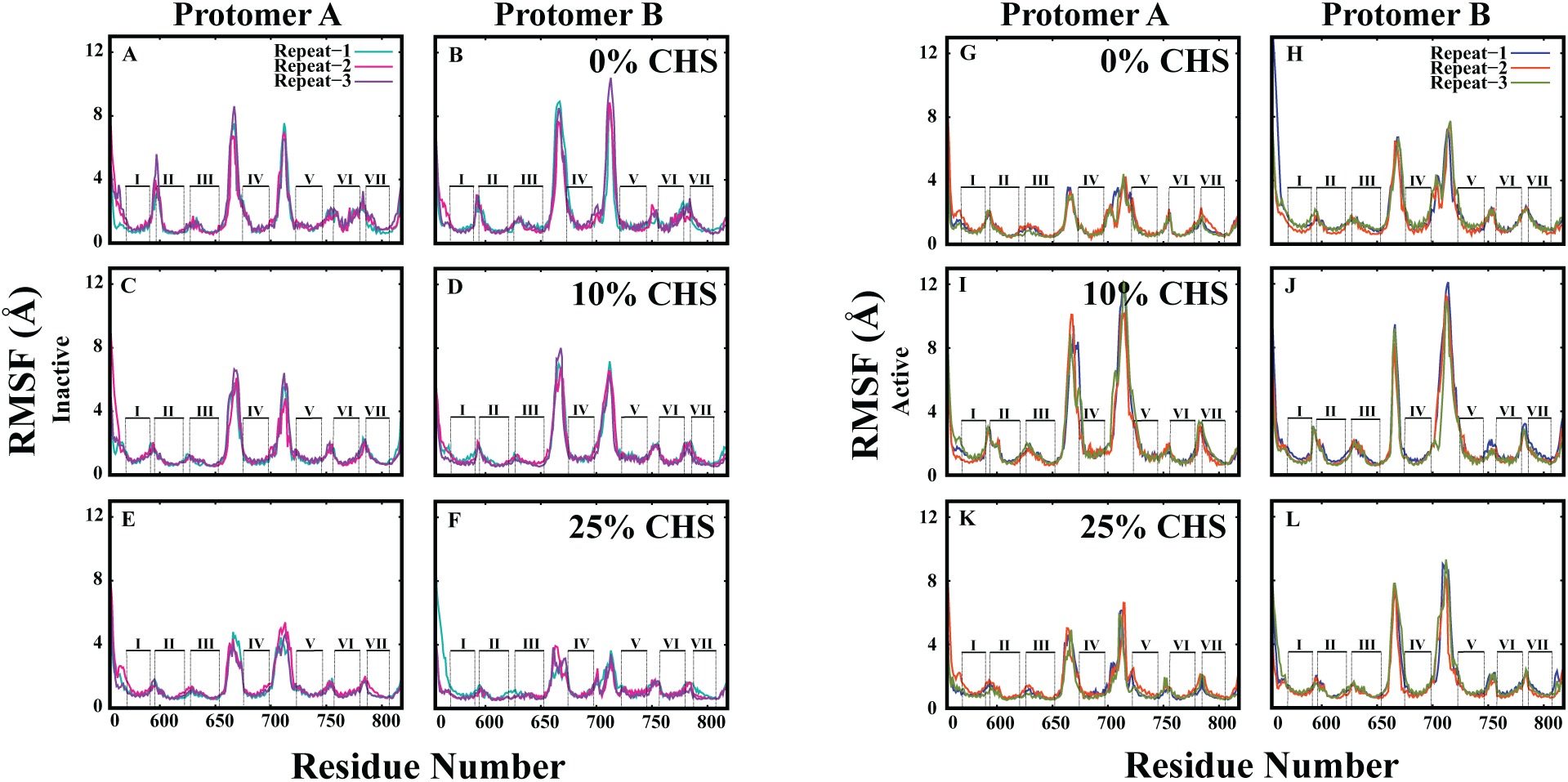
Root Mean Square Fluctuation (RMSF) analysis of mGluR2 embedded in a bilayer under different cholesteryl (CHS) concentrations. Panels on the left represent the RMSF profiles for Protomer A and Protomer B in the inactive state, while panels on the right depict the active state. Each panel shows the fluctuation of residues along the receptor sequence at 0%, 10%, and 25% CHS, with the seven transmembrane helices (I–VII) indicated. The three colored lines in each panel correspond to independent simulation replicates. Protomer A and Protomer B exhibit distinct fluctuation patterns, particularly in transmembrane helices IV and VI, where cholesteryl concentration modulates receptor flexibility.

In the inactive state, the loops between helices 1–2, 3–4, 4–5, and 6–7 showed the greatest flexibility at 0% CHS, decreasing progressively at 10% and 25% CHS. This mirrors the micelle observations and suggests that CHS reduces receptor flexibility by reinforcing interhelical packing in the inactive state.

However, in the active state, 10% CHS led to the highest loop flexibility in both protomers A and B, particularly within the same interhelical regions. In contrast, 0% and 25% CHS showed reduced fluctuations. This result diverges from the micelle environment, where CHS consistently increased flexibility in the active state. These differences highlight the influence of bilayer structure in modulating CHS effects and suggest that the bilayer allows 10% CHS to promote conformational transitions more effectively than in micelles.

Taken together, these findings suggest that CHS stabilizes the inactive state in both environments but exerts environment-specific effects in the active state. In bilayers, 10% CHS enhances flexibility, potentially enabling access to activation intermediates, while 25% CHS stabilizes a more rigid conformation. These observations underscore the importance of membrane composition in regulating GPCR dynamics and function.

### Cholesterol-Dependent Modulation of mGluR2 Dimer Stability and Signaling in a Bilayer

To investigate how the bilayer environment influences mGluR2 architecture, we analyzed inter-protomer distance and orientation across CHS concentrations (0%, 10%, and 25%) in both inactive and active states. While CHS has previously been shown to regulate dimeric stability in micelle systems, its effects within a lipid bilayer exhibit distinct patterns (Figure 11 and S11).

In the bilayer, the inter-protomer distance was highest at 10% CHS in both inactive and active states, in contrast to the monotonic increase seen in micelles. This suggests that intermediate CHS levels promote a more expanded dimeric conformation, potentially increasing conformational sampling and reducing structural constraints. At 25% CHS, mGluR2 adopted a more compact arrangement, likely reflecting a stabilized conformation.

Conversely, the inter-protomer angle exhibited a minimum at 10% CHS across both states. This inverse relationship indicates that while protomers are farther apart at 10% CHS, their relative orientation is more tightly constrained. Such a configuration may enable dynamic inter-protomer flexibility while preserving alignment for functional interactions (Figure 11 and S12). In contrast, both 0% and 25% CHS produced higher angular deviations, reflecting either increased structural fluctuation or excessive rigidity.

**Fig. 11.**
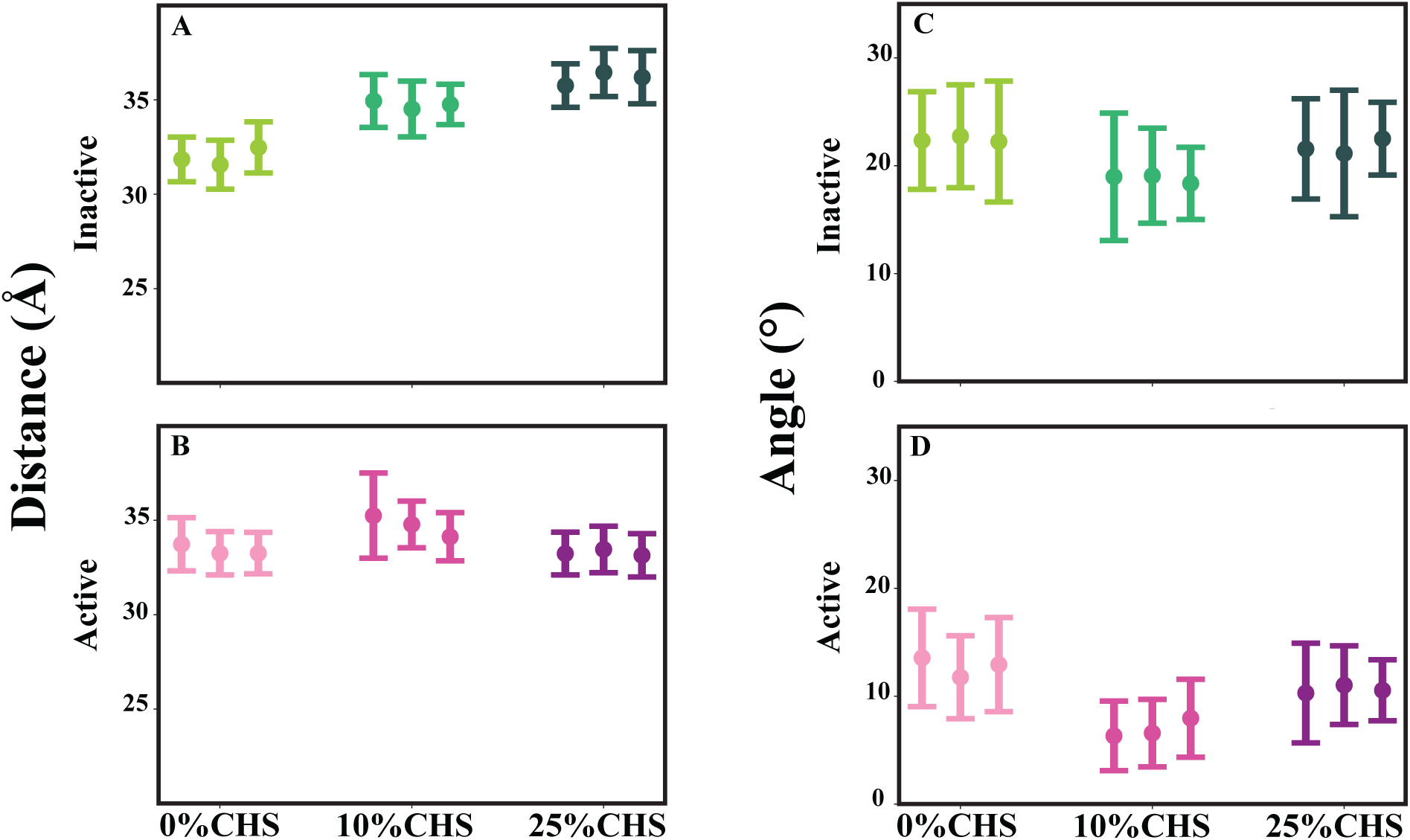
Interprotomer Distance and Angle Analysis of mGluR2 in Active and Inactive States within a bilayer. (A) Average interprotomer distance between Protomer A and Protomer B of the mGluR2 receptor in the inactive state under varying CHS concentrations (0%, 10%, and 25%), calculated from three independent simulations. (B) Average interprotomer distance for the active state under the same CHS conditions. (C) Average interprotomer angle between Protomer A and Protomer B in the inactive state, highlighting structural differences under varying CHS concentrations. (D) Average interprotomer angle for the active state. The error bars represent standard deviations, illustrating the impact of CHS concentrations on the structural stability and relative orientation of the two protomers in a bilayer environment.

These observations underscore how CHS modulates receptor architecture in a bilayer. At 10% CHS, the receptor adopts a unique balance of increased distance and constrained angle—potentially optimizing conditions for transitions between inactive and active states. Higher CHS levels (25%) appear to reinforce structural rigidity, possibly stabilizing a single active-like ensemble, while the CHS-free system displays greater angular variability, suggesting a more flexible but less coordinated state (Figure 11).

Overall, the bilayer imposes a distinct CHS-dependent modulation of dimeric geometry in mGluR2. These nonlinear trends highlight the critical role of lipid composition in shaping receptor activation dynamics and structural transitions, providing deeper insight into how cholesterol content may fine-tune receptor signaling in a membrane context.

### State-Dependent Structural Rearrangements of Individual TM Helices in mGluR2 Embedded in a Bilayer

Building on our findings from mGluR2 embedded in a micelle, we investigated how the bilayer environment influences state-dependent conformational dynamics of individual transmembrane (TM) helices across different CHS concentrations. As in the micelle system, CHS modulates helix stability and flexibility, but distinct trends emerge in the bilayer, suggesting that membrane composition plays a key role in fine-tuning receptor dynamics (Figure 12 and S13).

**Fig. 12.**
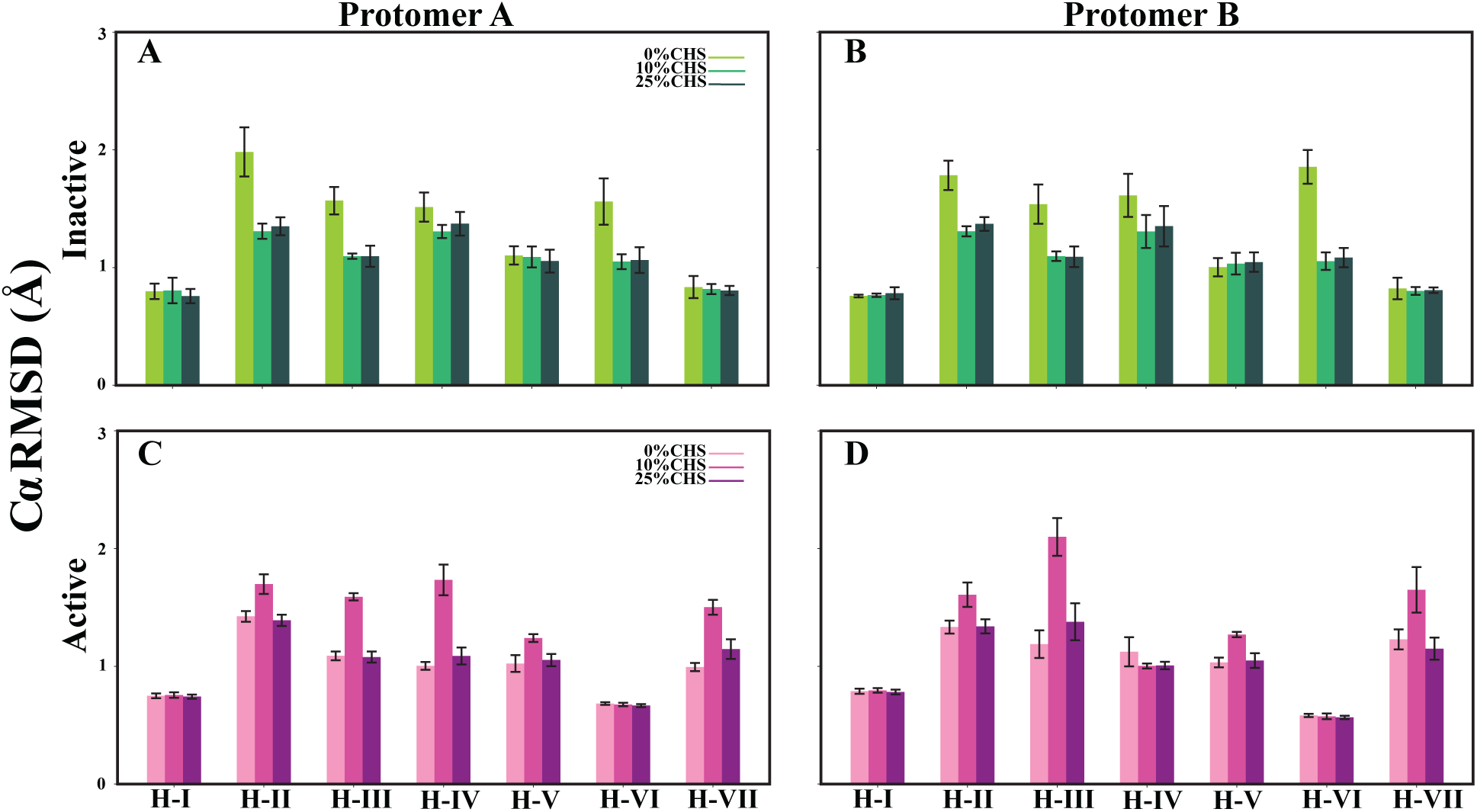
Mean and Standard Deviation of C*α* RMSD for Transmembrane Helices (H-I to H-VII) in mGluR2 Embedded in a Lipid Bilayer under Varying CHS Concentrations. RMSD data for individual helices are presented for Protomer A (panels A, C) and Protomer B (panels B, D) in both inactive (A–B) and active (C–D) states. Each bar represents the mean C*α* RMSD calculated over three independent simulation replicates, with error bars indicating standard deviation. Simulations were performed at three CHS concentrations (0%, 10%, and 25%) to examine how cholesterol modulates the structural flexibility of the transmembrane helices in a bilayer environment. The results reveal helix- and state-specific variations in stability, with certain helices—such as H-II and H-III in the active state—exhibiting enhanced RMSD at higher CHS levels, suggesting CHS-dependent conformational dynamics.

In the inactive state, TM1, TM5, and TM7 remained stable across all CHS conditions in both protomers A and B, consistent with their role as structural anchors. However, TM2, TM3, and TM6 displayed the highest RMSD values in the CHS-free system (0% CHS), indicating greater flexibility in the absence of CHS. This mirrors observations in the micelle system, where CHS-free conditions allowed greater structural fluctuations, possibly facilitating the receptor’s conformational transitions. Notably, TM6 exhibited the highest flexibility in both protomers at 0% CHS, reinforcing the idea that CHS stabilizes this helix and restricts its motion in the inactive state.

In the active state, TM1 and TM6 remained highly stable across all CHS conditions in both protomers, resembling their behavior in the inactive state. In contrast, TM2, TM3, TM5, and TM6 exhibited the highest RMSD values at 10% CHS, suggesting that this CHS concentration enhances flexibility in these helices, potentially facilitating activation-related conformational transitions. Interestingly, TM4 in protomer A also displayed the highest RMSD at 10% CHS, whereas in protomer B, it remained relatively stable across conditions. The increased flexibility at 10% CHS in the active state is a notable deviation from the micelle system, where higher CHS concentrations (25%) induced the most pronounced fluctuations (Figure 12 and S14). This suggests that in a bilayer, an intermediate CHS level may enhance receptor mobility in a way that supports structural rearrangements necessary for activation. Comparing the micelle and bilayer systems reveals key differences in CHS-dependent effects. In both environments, CHS-free conditions resulted in greater flexibility for TM2, TM3, and TM6 in the inactive state, reinforcing the role of CHS in stabilizing the receptor’s resting conformation. However, in the bilayer, 10% CHS introduced the most pronounced flexibility in the active state, particularly for TM2, TM3, TM5, and TM6, while in the micelle, higher CHS concentrations (25%) drove similar structural effects. This indicates that CHS acts as a modulator of GPCR dynamics differently in a lipid bilayer compared to a micelle, likely due to differences in lipid-protein interactions and membrane constraints.

From a functional perspective, these findings highlight the role of CHS in fine-tuning the conformational landscape of mGluR2. The stabilization of TM1 and TM6 across all conditions suggests a conserved structural framework, while increased flexibility at 10% CHS in the active state suggests a dynamic equilibrium that may facilitate receptor activation. In contrast, at higher CHS concentrations (25%), the receptor may adopt a more constrained active-like conformation, limiting its ability to explore intermediate states. The bilayer-dependent shift in CHS effects suggests that membrane composition plays a crucial role in receptor stability and activation dynamics, with CHS concentration acting as a key regulator of GPCR functional states.

### Cholesterol-Dependent Modulation of Hydrogen Bonding in mGluR2

Previously, we examined the influence of cholesteryl on hydrogen bond formation in mGluR2 embedded in a micelle, where the absence of a structured lipid bilayer allowed for increased flexibility and dynamic interactions between transmembrane (TM) helices. The micelle environment facilitated unique cholesteryl-dependent interactions, with key hydrogen bonds exhibiting asymmetric and state-dependent stability across receptor protomers. However, when embedded in a bilayer, mGluR2 is subjected to a more physiologically relevant lipid environment, where cholesteryl’s role in stabilizing or disrupting interhelical hydrogen bonds becomes more pronounced due to the structured nature of the membrane. To further explore how mGluR2 behaves in a bilayer, we analyzed the same key hydrogen bonds—ARG603–PHE589 (TM1–TM2), THR629–THR705 (TM3–loop (TM4–TM5)), and TRP773–SER801 (TM6–TM7)—to compare their state-dependent modulation in the presence of cholesteryl. Our findings indicate that while some trends observed in the micelle persist in the bilayer, new patterns emerge, reinforcing the idea that the lipid environment critically shapes receptor dynamics.

The TM1–TM2 interaction, mediated by the ARG603–ASN593 hydrogen bond, exhibited a distinct cholesteryl-dependent pattern. In the inactive state, this bond was absent in both Protomer A and Protomer B at 0% cholesteryl, suggesting an intrinsic instability in a cholesteryl-free bilayer, similar to what was observed in the micelle system. However, at 10% and 25% cholesteryl, the bond was consistently present in Protomer A, indicating that cholesteryl facilitates TM1–TM2 stabilization in the bilayer. In Protomer B at 10% cholesteryl, however, ARG603 did not engage in hydrogen bonding with ASN593, suggesting that alternative electrostatic interactions influenced the local stabilization of TM1–TM2 at this CHS concentration. This differentiates the bilayer response from the micelle system, where intermediate cholesteryl concentrations also influenced TM1–TM2 interactions, though via different mechanisms. In the active state, the hydrogen bond exhibited a different trend, forming at 0% and 25% cholesteryl in both protomers, while at 10% cholesteryl, it was disrupted (Figure 13). This behavior mirrors the micelle system, reinforcing the idea that intermediate cholesteryl levels induce a transient destabilization of TM1–TM2 interactions across different membrane environments. The absence of this bond in Protomer B at 10% CHS, along with the observed variability in electrostatic interactions, suggests that cholesteryl modulates TM1–TM2 stability in a state- and protomer-dependent manner.

**Fig. 13.**
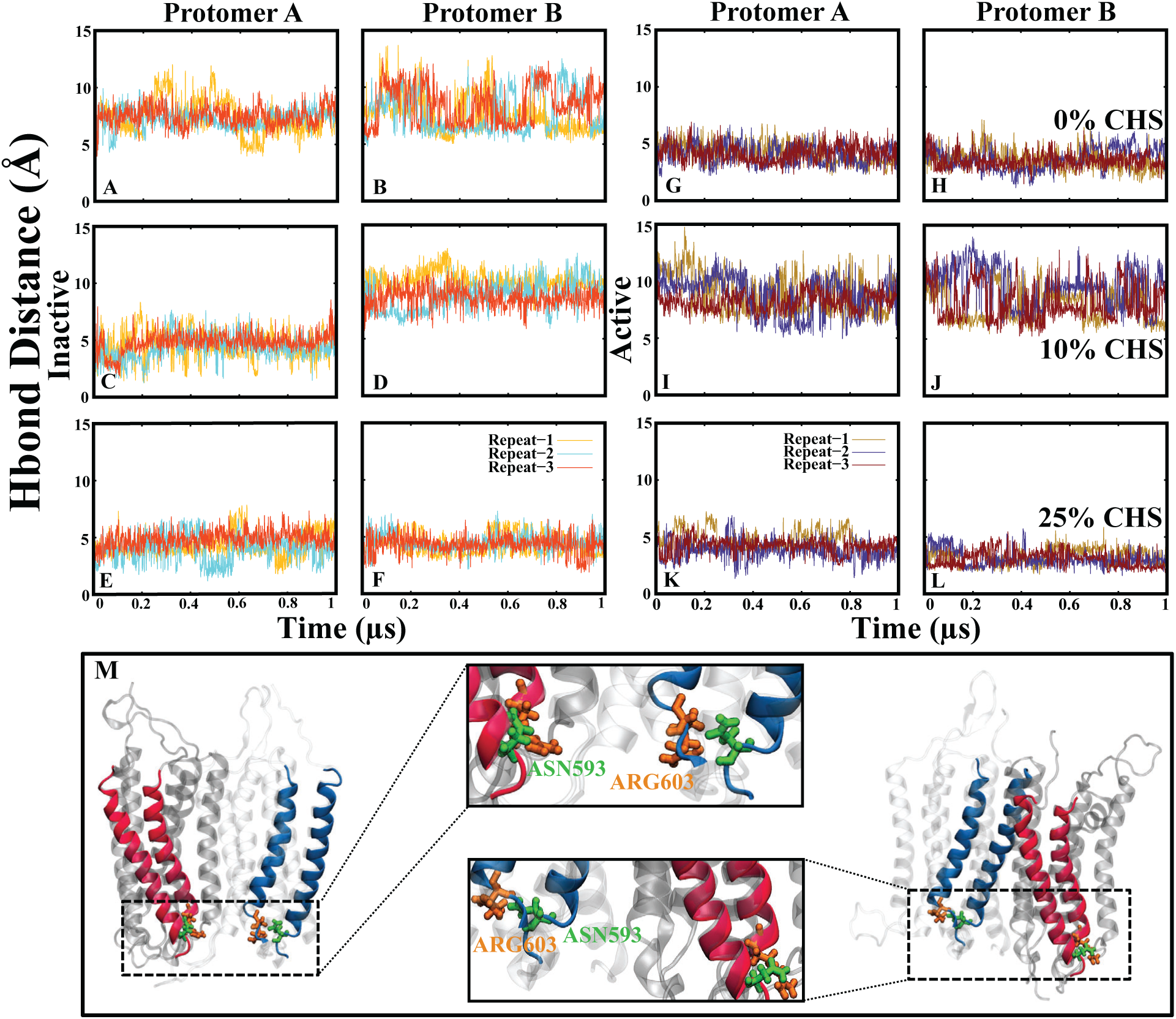
Hydrogen bond interaction between ASN593 and ARG603 stabilizes the TM1 and TM2 helices in the inactive and active states of mGluR2 embedded in a bilayer. Panels (A, B) show the time series of the hydrogen bond distances between the C*α* atoms of ARG603 (TM2) and ASN593 (TM1) at 0% CHS in Protomer A and Protomer B in the inactive state. Panels (C, D) and (E, F) display the corresponding distances at 10% CHS and 25% CHS in Protomer A and Protomer B in the inactive state, respectively. Panels (G, H) illustrate the hydrogen bond distances at 0% CHS in Protomer A and Protomer B in the active state, while panels (I, J) and (K, L) depict the data at 10% CHS and 25% CHS in Protomer A and Protomer B in the active state, respectively. Panel (M) presents a structural representation of the ARG603–ASN593 hydrogen bond interactions within the TM1–TM2 interface, highlighting their spatial positioning within the receptor’s transmembrane helices.

The TM3–loop (TM4–TM5) interaction, defined by the THR629–THR705 hydrogen bond, displayed a strong cholesteryl-dependent trend. In the inactive state, this bond was absent at 0% cholesteryl, similar to the micelle, indicating high TM3 flexibility in cholesteryl-free conditions. However, at 10% and 25% cholesteryl, the bond was consistently present, reinforcing the idea that cholesteryl enhances structural rigidity in both micelle and bilayer systems. In the active state, the bond was fully formed at 0% cholesteryl, whereas at 10% cholesteryl, it failed to form, mirroring the micelle system’s behavior. Interestingly, at 25% cholesteryl, the bond appeared midway through the simulation (Figure 14), suggesting a delayed stabilization effect that was less evident in the micelle, highlighting potential differences in cholesteryl’s role in a bilayer versus a micelle.

**Fig. 14.**
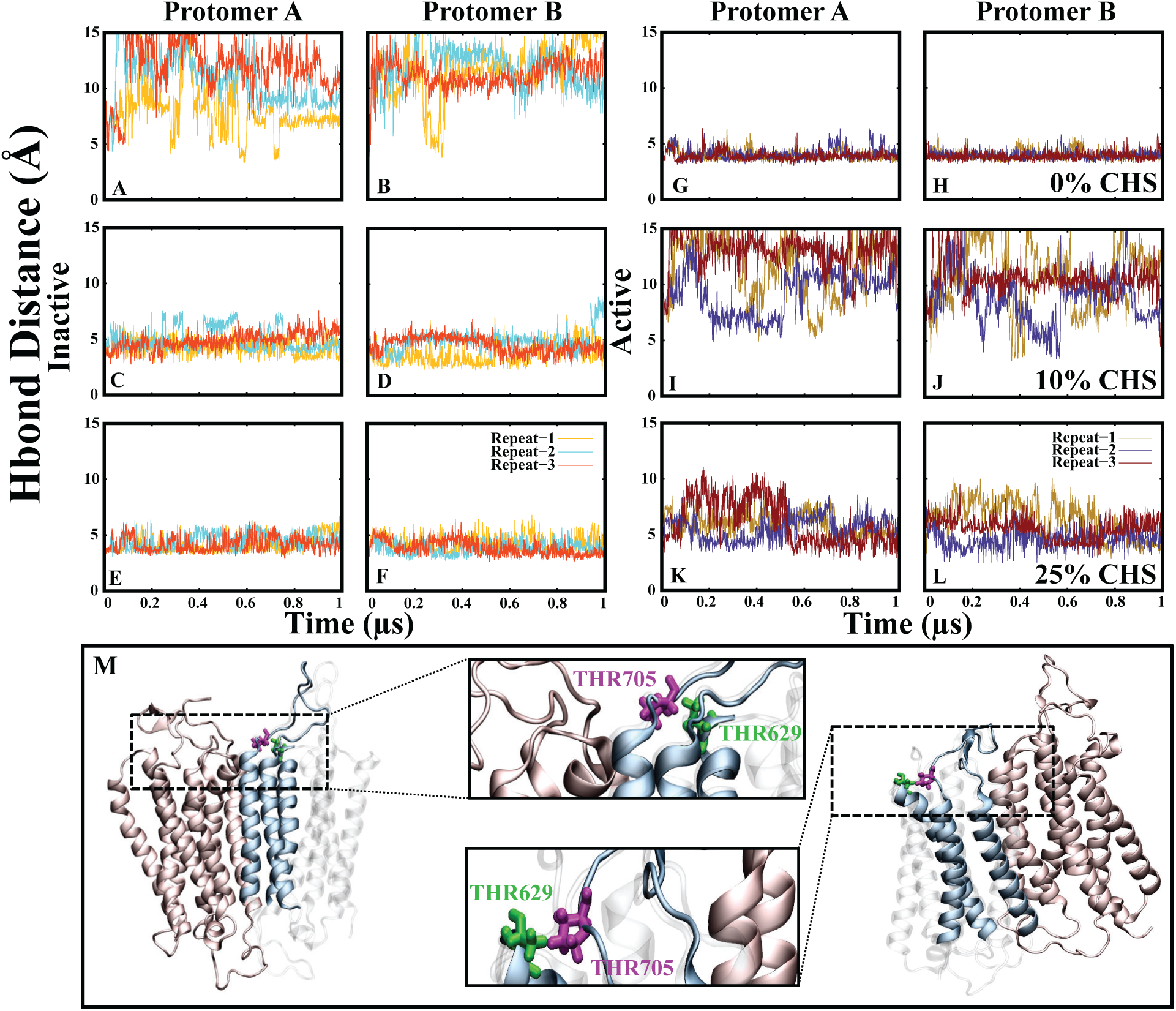
Hydrogen bond interaction between THR629 and THR705 stabilizes the TM3–loop (TM4–TM5) region in the inactive and active states of mGluR2 embedded in a bilayer. Panels (A, B) represent the time series of the hydrogen bond distances between the C*α* atoms of THR629 (TM3) and THR705 (loop between TM4–TM5) at 0% CHS in Protomer A and B in the inactive state. Panels (C, D) and (E, F) show the corresponding distances at 10% CHS and 25% CHS. Panels (G, H) illustrate the hydrogen bond distances at 0% CHS in the active state. Panels (I, J) and (K, L) depict the data at 10% and 25% CHS in the active state. Panel (M) provides a structural representation of the THR629–THR705 hydrogen bond.

The TM6–TM7 interaction, associated with the TRP773–SER801 hydrogen bond, also showed cholesteryl-sensitive behavior. In the inactive state, this bond was absent at 0% cholesteryl, similar to the micelle condition, indicating that TM6–TM7 flexibility is a shared feature in both environments under cholesteryl-free conditions. At 10% cholesteryl, the bond did not form immediately but appeared later in the simulation, showing a delayed stabilization effect from cholesteryl, just as in the micelle. However, at 25% cholesteryl, the bond was consistently present, contrasting with the micelle system, where excessive cholesteryl disrupted this interaction. This suggests that in a bilayer, cholesteryl promotes TM6–TM7 stability at higher concentrations rather than destabilizing it, possibly due to differences in lipid packing. In the active state, the bond was consistently formed at 0% and 25% cholesteryl, while at 10% cholesteryl (Figure 15), it was absent—mirroring the micelle trend, where intermediate cholesteryl levels appeared to disrupt TM6–TM7 stabilization.

**Fig. 15.**
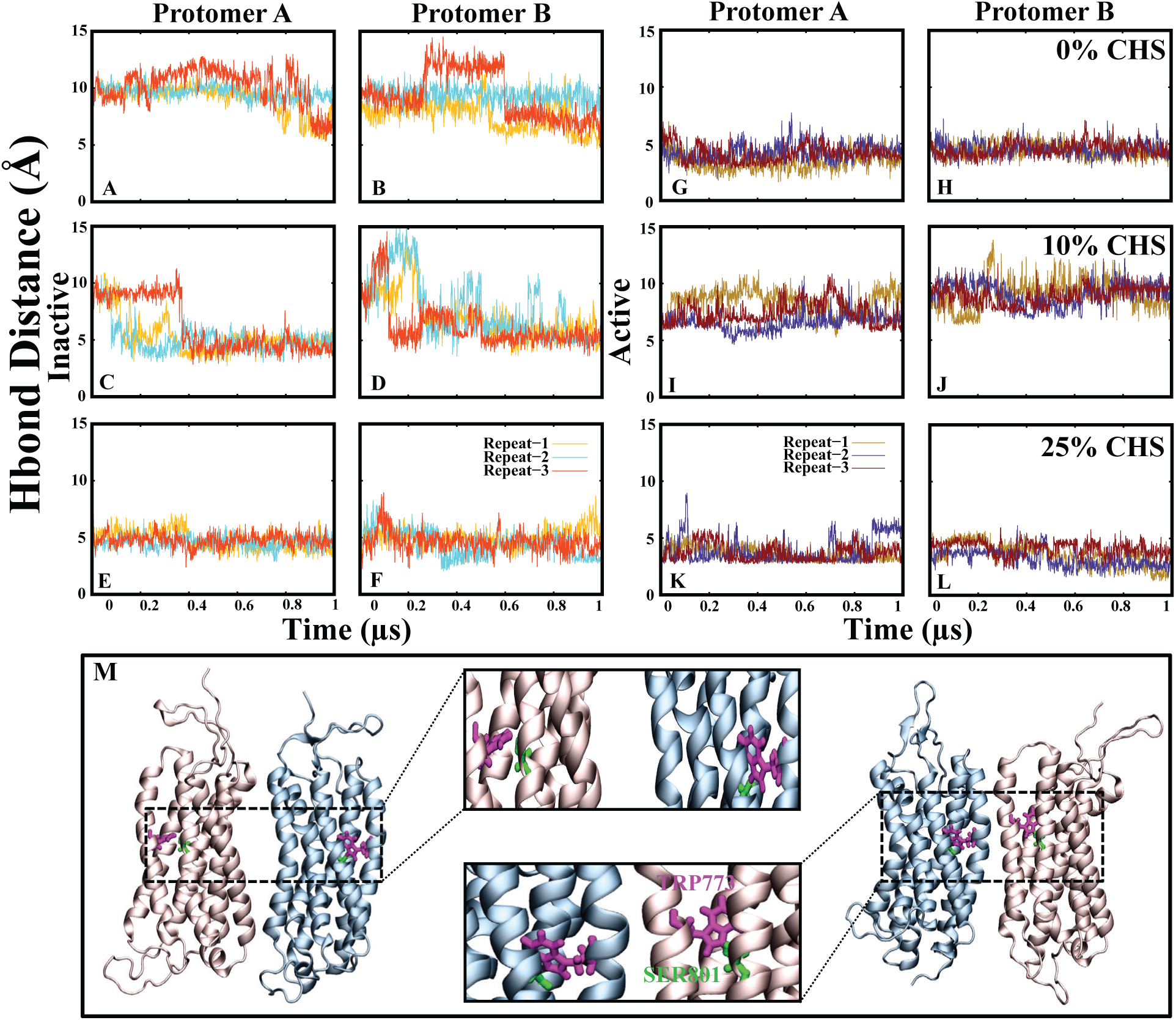
Hydrogen bond interaction between TRP773 and SER801 stabilizes the TM6 and TM7 helices in the inactive and active states of mGluR2 embedded in a bilayer. Panels (A, B) represent the time series of the hydrogen bond distances between the C*α* atoms of TRP773 (TM6) and SER801 (TM7) at 0% CHS in Protomer A and Protomer B in the inactive state. Panels (C, D) and (E, F) show the corresponding distances at 10% CHS and 25% CHS in Protomer A and Protomer B in the inactive state, respectively. Panels (G, H) illustrate the hydrogen bond distances at 0% CHS in Protomer A and Protomer B in the active state, while panels (I, J) and (K, L) depict the data at 10% CHS and 25% CHS in Protomer A and Protomer B in the active state, respectively. Panel (M) provides a structural representation of the TRP773–SER801 hydrogen bond interactions within the TM6–TM7 interface, highlighting their positioning within the receptor’s transmembrane helices.

By comparing the micelle and bilayer environments, it is evident that while certain cholesteryl-dependent hydrogen bonding trends persist across both systems, the bilayer provides additional structural constraints that influence receptor behavior. The presence of an ordered lipid membrane enhances cholesteryl’s stabilizing effects at higher concentrations, whereas in a micelle, cholesteryl’s influence appears more transient and asymmetric. These findings emphasize the importance of lipid composition in shaping mGluR2 function and suggest that cholesteryl’s role in allosteric regulation is context-dependent, varying between simplified micelle environments and more complex bilayer structures.

### Salt Bridge Dynamics and Their Relationship to Protomer Distance and Angle in mGluR2 Embedded in a Bilayer

Previous studies of mGluR2 embedded in a micelle revealed a cholesteryl (CHS)-dependent network of salt bridges that played a key role in stabilizing receptor conformations, modulating inter-protomer distance, and influencing angle fluctuations. However, when mGluR2 is embedded in a bilayer, the structured lipid environment imposes additional constraints on receptor flexibility, altering the CHS-induced modulation of salt bridge interactions. By examining the formation and stability of key inter-protomer salt bridges, we observed distinct patterns in the bilayer (Figure 16), some of which align with trends seen in the micelle, while others reveal bilayer-specific effects on receptor dynamics.

**Fig. 16.**
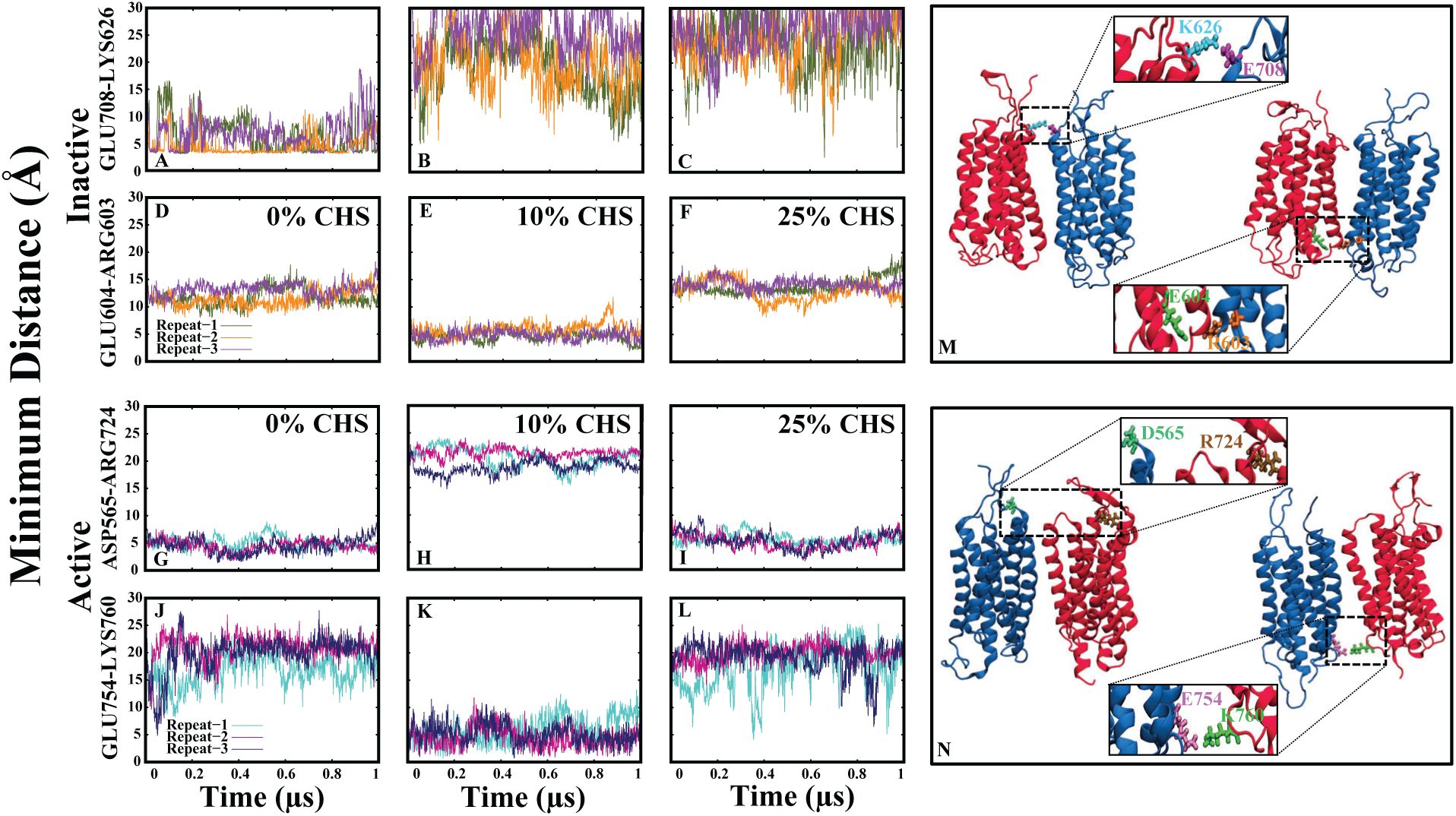
Minimum distance analysis of salt-bridge interactions in mGluR2 embedded in a bilayer. Panels (A, B, C) show the time series of the minimum distances between Glu708-Lys626 in the inactive state at 0%, 10%, and 25% CHS, respectively. Panels (D, E, F) display the minimum distances of the Glu604-Arg603 interaction in the inactive state at 0%, 10%, and 25% CHS. Panels (G, H, I) represent the minimum distances of the Asp565-Arg724 interaction in the active state at 0%, 10%, and 25% CHS. Panels (J, K, L) show the minimum distance analysis of the Glu754-Lys760 interaction in the active state at 0%, 10%, and 25% CHS. The three colored lines in each panel correspond to independent simulation replicates.

In the inactive state, a salt bridge between Glu708 (Helix 4, Protomer B) and Lys626 (Helix 2, Protomer A) was consistently formed at 0% and 25% cholesteryl hemisuccinate (CHS), stabilizing a more compact receptor conformation under these conditions. In contrast, at 10% CHS, this interaction was absent, and the inter-protomer distance was significantly higher, indicating that an intermediate CHS concentration disrupts this electrostatic interaction, promoting a more expanded receptor conformation. A comparable effect has been observed in simulations of the *β*_2_ adrenergic receptor, where the loss of the ionic lock (Arg131(3.50)–Glu268(6.30)) in a POPC/cholesterol bilayer resulted in increased receptor flexibility and destabilization of the inactive state.^21^ This effect resembles CHS-mediated disruptions observed in the micelle, where intermediate CHS levels also altered inter-helical interactions, likely due to partial stabilization of cholesterol-binding sites that interfere with salt bridge formation.

Additionally, in the inactive state, another key inter-protomer salt bridge between Glu604 (Helix 2, Protomer A) and Arg603 (Helix 2, Protomer B) was formed at 10% CHS, correlating with a lower inter-protomer angle at this concentration. However, a hydrogen bond that was expected to form in Protomer B at 10% CHS was absent, likely due to Arg603 in Protomer B forming a salt bridge with Glu604 in Protomer A instead. A similar salt bridge rearrangement has been observed in the calcium-sensing receptor (CaSR), where the R680G mutation disrupted the Arg680–Glu767 interaction, shifting receptor stabilization mechanisms and leading to signaling bias through *β*-arrestin instead of G-proteins.^71^ This shift in interaction patterns suggests that CHS fine-tunes inter-protomer electrostatic contacts, leading to alternative stabilization mechanisms at different CHS concentrations.

In the active state, a salt bridge between Asp656 (Helix 1, Protomer A) and Arg724 (Helix 5, Protomer B) was present at 0% and 25% CHS, indicating that at these concentrations, electrostatic interactions contribute to receptor activation. However, at 10% CHS, this salt bridge was absent, and the inter-protomer distance was higher, suggesting that an intermediate CHS concentration disrupts this stabilizing interaction, mirroring trends seen in the inactive state. This is consistent with findings on the glucagon receptor (GCGR), where activation requires overcoming a structural energy barrier, and the outward movement of TM6 is only triggered by G protein engagement rather than agonist binding alone.^72^

Another significant active-state salt bridge was observed between Glu754 (Helix 5, Protomer A) and Lys760 (Helix 6, Protomer B) at 10% CHS, which correlated with a lower inter-protomer angle at this concentration. This suggests that while 10% CHS disrupts certain stabilizing salt bridges, such as Asp656-Arg724, it simultaneously facilitates alternative inter-helical interactions. This aligns with findings in the gonadotropin-releasing hormone receptor (GnRHR), where critical salt bridges (Arg38-Asp98 and Glu90-Lys121) were shown to regulate receptor trafficking and activation.^73^ Similarly, this aligns with findings in the cannabinoid receptor 1 (CB1), where disruption of key salt bridges (R2.37-D6.30 and D2.63-K3.28) was found to drive transitions between inactive and active states.^66^ In particular, breaking the R2.37-D6.30 interaction resulted in enhanced G-protein coupling and increased receptor flexibility, consistent with our observation that CHS-dependent salt bridge disruption promotes receptor conformational shifts.

These findings highlight a state- and CHS-dependent salt bridge network that influences inter-protomer distance, angle fluctuations, and receptor stability in mGluR2 embedded in a bilayer. While some trends observed in the micelle persist in the bilayer, the structured environment of the bilayer imposes additional constraints, leading to distinct CHS-dependent effects on receptor stability and activation. Low CHS (0%) and high CHS (25%) concentrations favor compact receptor conformations, stabilizing key salt bridges such as Glu708-Lys626 in the inactive state and Asp656-Arg724 in the active state. Intermediate CHS levels (10%) disrupt certain stabilizing interactions, increasing inter-protomer distance, but also facilitate alternative salt bridge formation, such as Glu754-Lys760 in the active state. CHS concentration fine-tunes receptor stability by selectively modulating inter-protomer electrostatic interactions, influencing the receptor’s conformational equilibrium and activation efficiency. A comparison between bilayer-embedded and micelle-embedded mGluR2 suggests that CHS-induced salt bridge rearrangements are strongly influenced by membrane composition and receptor environment. These results provide insights into how lipid bilayers regulate receptor structure and function, with potential implications for membrane-protein interactions and CHS-mediated allosteric modulation of GPCRs.

## Conclusion

Our study provides a detailed examination of the lipid-mediated modulation of mGluR2 in both micelle and bilayer environments using all-atom molecular dynamics simulations. By analyzing the effects of varying cholesteryl hemisuccinate (CHS) concentrations (0%, 10%, and 25%), we elucidate key structural and dynamic changes that contribute to receptor stability, inter-protomer interactions, and functional behavior.

Our results reveal a state-dependent effect of CHS on receptor stability. In the micelle environment, 0% CHS led to the highest flexibility in the inactive state, allowing for greater conformational sampling, while 25% CHS stabilized a more compact receptor conformation, particularly in the active state. The intermediate CHS concentration (10%) exhibited the most pronounced conformational transitions, indicating a possible balance between receptor flexibility and structural stability that may be critical for activation.

At the transmembrane helix (TM) level, TM1 remained stable across all conditions, reinforcing its role as a structural anchor. However, TM3, TM4, and TM5 exhibited notable flexibility, particularly in the absence of CHS, suggesting a cholesterol-dependent modulation of key structural elements involved in receptor activation. In contrast, TM6 and TM7 displayed increased fluctuations at 25% CHS, potentially stabilizing the active-like state while restricting the receptor’s ability to transition to alternate conformations. The asymmetric behavior observed between protomers further suggests a cholesterol-driven influence on receptor dimerization and functional transitions.

Our hydrogen bond and salt bridge analyses provide further insights into how CHS concentration modulates key intra- and inter-helical interactions. Higher CHS concentrations promoted stronger inter-helical contacts, reducing the overall receptor flexibility, whereas lower CHS concentrations facilitated a more dynamic conformational landscape. Notably, the inter-protomer distance and orientation fluctuated in a CHS-dependent manner, with the 10% CHS system demonstrating a unique compaction pattern that may facilitate receptor activation.

In the bilayer environment, CHS had a differential impact on receptor stability and flexibility. The active state exhibited increased conformational fluctuations at 10% CHS, while 25% CHS enforced a more rigid conformation. The loop regions connecting TM helices showed a CHS-dependent flexibility pattern, with increased mobility at 10% CHS, suggesting a potential role in facilitating functional transitions. These findings indicate that while CHS stabilizes specific conformations, an intermediate concentration allows for a dynamic equilibrium essential for receptor activation.

Taken together, our findings reinforce the critical role of CHS in modulating mGluR2 structure and function. The differential effects of CHS concentration on receptor stability, flexibility, and inter-protomer interactions highlight the importance of lipid composition in GPCR regulation. These insights contribute to our broader understanding of GPCR-lipid interactions and offer valuable implications for drug discovery efforts targeting lipid-mediated receptor modulation. Future studies should explore the impact of additional lipid components and extend these simulations to longer timescales to further refine our understanding of cholesterol-governed receptor function.

## Supporting Information Available

### Data availability

https://github.com/bslgroup/mglur2.git

## Acknowledgement

This research was supported by the National Institute of General Medical Sciences (NIH grant R35GM147423 awarded to M.M.), the National Science Foundation (NSF grant CHE 1945465 awarded to M.M.), and the Arkansas Biosciences Institute. Computational resources were provided by the Texas Advanced Computing Center (TACC) at the University of Texas at Austin (Frontera) through LRAC allocation CHE21003 to M.M. The work also used Stampede at TACC, Expanse at the San Diego Supercomputer Center, and Bridges-2 at the Pittsburgh Supercomputing Center through allocation MCB150129 from the Advanced Cyberinfrastructure Coordination Ecosystem: Services & Support (ACCESS) program, supported by NSF grants #2138259, #2138286, #2138307, #2137603, and #2138296.^74^ Additional computational support came from the Arkansas High-Performance Computing Center, funded by multiple NSF grants and the Arkansas Economic Development Commission.

## Notes

### Competing Interest Statement

The authors have declared no competing interest.

## References

(1) Yang, D.; Zhou, Q.; Labroska, V.; Qin, S.; Darbalaei, S.; Wu, Y.; Yuliantie, E.; Xie, L.; Tao, H.; Cheng, J., et al. G protein-coupled receptors: structure-and function-based drug discovery. Signal transduction and targeted therapy 2021, 6, 7.

(2) Cheng, L.; Xia, F.; Li, Z.; Shen, C.; Yang, Z.; Hou, H.; Sun, S.; Feng, Y.; Yong, X.; Tian, X. et al. Structure, function and drug discovery of GPCR signaling. Molecular Biomedicine 2023, 4, 46.

(3) Arang, N.; Gutkind, J. S. G Protein-Coupled receptors and heterotrimeric G proteins as cancer drivers. FEBS letters 2020, 594, 4201–4232.

(4) Crupi, R.; Impellizzeri, D.; Cuzzocrea, S. Role of metabotropic glutamate receptors in neurological disorders. Frontiers in molecular neuroscience 2019, 12, 20.

(5) Srivastava, A.; Das, B.; Yao, A. Y.; Yan, R. Metabotropic glutamate receptors in Alzheimer’s disease synaptic dysfunction: Therapeutic opportunities and hope for the future. Journal of Alzheimer’s Disease 2020, 78, 1345–1361.

(6) Wong, T.-S.; Li, G.; Li, S.; Gao, W.; Chen, G.; Gan, S.; Zhang, M.; Li, H.; Wu, S.; Du, Y. G protein-coupled receptors in neurodegenerative diseases and psychiatric disorders. Signal Transduction and Targeted Therapy 2023, 8, 177.

(7) Jin, L. E.; Wang, M.; Yang, S.-T.; Yang, Y.; Galvin, V. C.; Lightbourne, T. C.; Ottenheimer, D.; Zhong, Q.; Stein, J.; Raja, A. et al. mGluR2/3 mechanisms in primate dorsolateral prefrontal cortex: evidence for both presynaptic and postsynaptic actions. Molecular psychiatry 2017, 22, 1615–1625.

(8) Dogra, S.; Conn, P. J. Targeting metabotropic glutamate receptors for the treatment of depression and other stress-related disorders. Neuropharmacology 2021, 196, 108687.

(9) Sweeten, B. L.; Adkins, A. M.; Wellman, L. L.; Sanford, L. D. Group II metabotropic glutamate receptor activation in the basolateral amygdala mediates individual differences in stress-induced changes in rapid eye movement sleep. Progress in Neuro-Psychopharmacology and Biological Psychiatry 2021, 104, 110014.

(10) Wolf, D. H.; Zheng, D.; Kohler, C.; Turetsky, B. I.; Ruparel, K.; Satterthwaite, T. D.; Elliott, M. A.; March, M. E.; Cross, A. J.; Smith, M. A. et al. Effect of mGluR2 positive allosteric modulation on frontostriatal working memory activation in schizophrenia. Molecular psychiatry 2022, 27, 1226–1232.

(11) Huang, L.; Xiao, W.; Wang, Y.; Li, J.; Gong, J.; Tu, E.; Long, L.; Xiao, B.; Yan, X.; Wan, L. Metabotropic glutamate receptors (mGluRs) in epileptogenesis: an update on abnormal mGluRs signaling and its therapeutic implications. Neural Regeneration Research 2024, 19, 360–368.

(12) Lin, S.; Han, S.; Cai, X.; Tan, Q.; Zhou, K.; Wang, D.; Wang, X.; Du, J.; Yi, C.; Chu, X. et al. Structures of Gi-bound metabotropic glutamate receptors mGlu2 and mGlu4. Nature 2021, 594, 583–588.

(13) Huang, S.; Cao, J.; Jiang, M.; Labesse, G.; Liu, J.; Pin, J.-P.; Rondard, P. Interdomain movements in metabotropic glutamate receptor activation. Proceedings of the National Academy of Sciences 2011, 108, 15480–15485.

(14) Song, W.; Yen, H.-Y.; Robinson, C. V.; Sansom, M. S. State-dependent lipid interactions with the A2a receptor revealed by MD simulations using in vivo-mimetic membranes. Structure 2019, 27, 392–403.

(15) Ray, A. P.; Jin, B.; Eddy, M. T. The conformational equilibria of a human GPCR compared between lipid vesicles and aqueous solutions by integrative 19F-NMR. bioRxiv 2024,

(16) Kurth, M.; Lolicato, F.; Sandoval-Perez, A.; Amaya-Espinosa, H.; Teslenko, A.; Sinning, I.; Beck, R.; Brügger, B.; Aponte-Santamaría, C. Cholesterol localization around the metabotropic glutamate receptor 2. The Journal of Physical Chemistry B 2020, 124, 9061–9078.

(17) Serdiuk, T.; Manna, M.; Zhang, C.; Mari, S. A.; Kulig, W.; Pluhackova, K.; Kobilka, B. K.; Vattulainen, I.; Müller, D. J. A cholesterol analog stabilizes the human *β*2-adrenergic receptor nonlinearly with temperature. Science signaling 2022, 15, eabi7031.

(18) Vukoti, K.; Kimura, T.; Macke, L.; Gawrisch, K.; Yeliseev, A. Stabilization of functional recombinant cannabinoid receptor CB2 in detergent micelles and lipid bilayers. 2012,

(19) Augustyn, B.; Stepien, P.; Poojari, C.; Mobarak, E.; Polit, A.; Wisniewska-Becker, A.; Róg, T. Cholesteryl hemisuccinate is not a good replacement for cholesterol in lipid nanodiscs. The Journal of Physical Chemistry B 2019, 123, 9839–9845.

(20) Tran, N. Lipid environment determines the drug-stimulated ATPase activity of P-glycoprotein. Biophysical Journal 2023, 122, 209a.

(21) Mahmood, M. I.; Liu, X.; Neya, S.; Hoshino, T. Influence of lipid composition on the structural stability of g-protein coupled receptor. Chemical and Pharmaceutical Bulletin 2013, 61, 426–437.

(22) Isu, U. H.; Badiee, S. A.; Polasa, A.; Tabari, S. H.; Derakhshani-Molayousefi, M.; Moradi, M. Cholesterol Dependence on the Conformational Changes of Metabotropic Glutamate Receptor 1. bioRxiv 2024, 2024–04.

(23) Isu, U. H.; Polasa, A.; Moradi, M. Differential Behavior of Conformational Dynamics in Active and Inactive States of Cannabinoid Receptor 1. The Journal of Physical Chemistry B 2024, 128, 8437–8447.

(24) Isu, U. H.; Badiee, S. A.; Khodadadi, E.; Moradi, M. Cholesterol in Class C GPCRs: Role, relevance, and localization. Membranes 2023, 13, 301.

(25) Venkatakrishnan, A.; others GPCR dynamics: structures in motion. 2017,

(26) Bhattarai, A.; Wang, J.; Miao, Y. G-protein-coupled receptor–membrane interactions depend on the receptor activation state. Journal of computational chemistry 2020, 41, 460–471.

(27) Torrens-Fontanals, M.; Stepniewski, T. M.; Aranda-García, D.; Morales-Pastor, A.; Medel-Lacruz, B.; Selent, J. How do molecular dynamics data complement static structural data of GPCRs. International journal of molecular sciences 2020, 21, 5933.

(28) Bruzzese, A.; Dalton, J. A.; Giraldo, J. Insights into adenosine A2A receptor activation through cooperative modulation of agonist and allosteric lipid interactions. PLoS computational biology 2020, 16, e1007818.

(29) Fleetwood, O.; Matricon, P.; Carlsson, J.; Delemotte, L. Energy landscapes reveal agonist control of G protein-coupled receptor activation via microswitches. Biochemistry 2020, 59, 880–891.

(30) Do, H. N.; Akhter, S.; Miao, Y. Pathways and mechanism of caffeine binding to human adenosine A2A receptor. Frontiers in molecular biosciences 2021, 8, 673170.

(31) Jo, J.; Heon, S.; Kim, M.; Son, G.; Park, Y.; Henley, J.; Weiss, J.; Sheng, M.; Collingridge, G.; Cho, K. Metabotropic Glutamate Receptor-mediated LTD Involves two Interacting Ca2+ Sensors, NCS-1 and PICK1. European Psychiatry 2009, 24, 24– E84.

(32) Wu, E. L.; Cheng, X.; Jo, S.; Rui, H.; Song, K. C.; Dávila-Contreras, E. M.; Qi, Y.; Lee, J.; Monje-Galvan, V.; Venable, R. M., et al. CHARMM-GUI membrane builder toward realistic biological membrane simulations. 2014.

(33) Lee, J.; Cheng, X.; Jo, S.; MacKerell, A. D.; Klauda, J. B.; Im, W. CHARMM-GUI input generator for NAMD, GROMACS, AMBER, OpenMM, and CHARMM/OpenMM simulations using the CHARMM36 additive force field. Biophysical journal 2016, 110, 641a.

(34) Cheng, X.; Kim, J.-K.; Kim, Y.; Bowie, J. U.; Im, W. Molecular dynamics simulation strategies for protein–micelle complexes. Biochimica et Biophysica Acta (BBA)- Biomembranes 2016, 1858, 1566–1572.

(35) Krüger, D. M.; Kamerlin, S. C. Micelle Maker: An online tool for generating equilibrated micelles as direct input for molecular dynamics simulations. ACS omega 2017, 2, 4524–4530.

(36) Best, R. B.; Zhu, X.; Shim, J.; Lopes, P. E.; Mittal, J.; Feig, M.; MacKerell Jr, A. D. Optimization of the additive CHARMM all-atom protein force field targeting improved sampling of the backbone *φ*, *ψ* and side-chain *χ*1 and *χ*2 dihedral angles. Journal of chemical theory and computation 2012, 8, 3257–3273.

(37) Klauda, J. B.; Venable, R. M.; Freites, J. A.; O’Connor, J. W.; Tobias, D. J.; Mondragon-Ramirez, C.; Vorobyov, I.; MacKerell Jr, A. D.; Pastor, R. W. Update of the CHARMM all-atom additive force field for lipids: validation on six lipid types. The journal of physical chemistry B 2010, 114, 7830–7843.

(38) Phillips, J. C.; Braun, R.; Wang, W.; Gumbart, J.; Tajkhorshid, E.; Villa, E.; Chipot, C.; Skeel, R. D.; Kale, L.; Schulten, K. Scalable molecular dynamics with NAMD. Journal of computational chemistry 2005, 26, 1781–1802.

(39) Harkey, T.; Govind Kumar, V.; Hettige, J.; Tabari, S. H.; Immadisetty, K.; Moradi, M. The role of a crystallographically unresolved cytoplasmic loop in stabilizing the bacterial membrane insertase yidc2. Scientific reports 2019, 9, 14451.

(40) Martyna, G. J.; Tobias, D. J.; Klein, M. L. Constant pressure molecular dynamics algorithms. The Journal of chemical physics 1994, 101, 4177–4189.

(41) Jo, S.; Kim, T.; Im, W. Automated builder and database of protein/membrane complexes for molecular dynamics simulations. PloS one 2007, 2, e880.

(42) Darden, T.; York, D.; Pedersen, L.; others Particle mesh Ewald: An N log (N) method for Ewald sums in large systems. Journal of chemical physics 1993, 98, 10089–10089.

(43) Humphrey, W.; Dalke, A.; Schulten, K. VMD: visual molecular dynamics. Journal of molecular graphics 1996, 14, 33–38.

(44) Illinois, U. Introduction to Molecular Dynamics VMD: Visual Molecular Dynamics. Biophysics 1996,

(45) Wu, Y.; Li, X.; Hua, T.; Liu, Z.-J.; Liu, H.; Zhao, S. MD simulations revealing special activation mechanism of cannabinoid receptor 1. Frontiers in Molecular Biosciences 2022, 9, 860035.

(46) Ji, S.; Yang, W.; Yu, W. Understanding the role of the CB1 toggle switch in interaction networks using molecular dynamics simulation. Scientific reports 2021, 11, 22369.

(47) Yokoi, S.; Mitsutake, A. Molecular dynamics simulations for the determination of the characteristic structural differences between inactive and active states of wild type and mutants of the orexin2 receptor. The Journal of Physical Chemistry B 2021, 125, 4286–4298.

(48) Liauw, B. W.-H.; Foroutan, A.; Schamber, M. R.; Lu, W.; Afsari, H. S.; Vafabakhsh, R. Conformational fingerprinting of allosteric modulators in metabotropic glutamate receptor 2. Elife 2022, 11, e78982.

(49) Xie, P.; Zhang, J.; Chen, B.; Li, X.; Zhang, W.; Zhu, M.; Li, W.; Li, J.; Fu, W. Computational Methods for Understanding the Selectivity and Signal Transduction Mechanism of Aminomethyl Tetrahydronaphthalene to Opioid Receptors. Molecules 2022, 27, 2173.

(50) Kolinski, M.; Filipek, S. Molecular dynamics of *μ* opioid receptor complexes with agonists and antagonists. Open Struct. Biol. J 2008, 2, 8–20.

(51) Bokoch, M. P.; Zou, Y.; Rasmussen, S. G.; Liu, C. W.; Nygaard, R.; Rosenbaum, D. M.; Fung, J. J.; Choi, H.-J.; Thian, F. S.; Kobilka, T. S. et al. Ligand-specific regulation of the extracellular surface of a G-protein-coupled receptor. Nature 2010, 463, 108–112.

(52) Liu, J. J.; Horst, R.; Katritch, V.; Stevens, R. C.; Wüthrich, K. Biased signaling pathways in *β*2-adrenergic receptor characterized by 19F-NMR. Science 2012, 335, 1106–1110.

(53) Thompson, A. A.; Liu, J. J.; Chun, E.; Wacker, D.; Wu, H.; Cherezov, V.; Stevens, R. C. GPCR stabilization using the bicelle-like architecture of mixed sterol-detergent micelles. Methods 2011, 55, 310–317.

(54) Banerjee, C.; Liauw, B. W.-H.; Vafabakhsh, R. Direct effect of membrane environment on the activation of mGluR2 revealed by single-molecule FRET. Structure 2025,

(55) Kobilka, B. K. G protein coupled receptor structure and activation. Biochimica et Biophysica Acta (BBA)-Biomembranes 2007, 1768, 794–807.

(56) Heyder, N. A.; Kleinau, G.; Speck, D.; Schmidt, A.; Paisdzior, S.; Szczepek, M.; Bauer, B.; Koch, A.; Gallandi, M.; Kwiatkowski, D. et al. Structures of active melanocortin-4 receptor–Gs-protein complexes with NDP-*α*-MSH and setmelanotide. Cell Research 2021, 31, 1176–1189.

(57) Farinha, A.; Lavreysen, H.; Peeters, L.; Russo, B.; Masure, S.; Trabanco, A.; Cid, J.; Tresadern, G. Molecular determinants of positive allosteric modulation of the human metabotropic glutamate receptor 2. British journal of pharmacology 2015, 172, 2383–2396.

(58) Díaz, Ó.; Dalton, J. A.; Giraldo, J. Revealing the mechanism of agonist-mediated cannabinoid receptor 1 (CB1) activation and phospholipid-mediated allosteric modulation. Journal of Medicinal Chemistry 2019, 62, 5638–5654.

(59) Manna, M.; Niemelä, M.; Tynkkynen, J.; Javanainen, M.; Kulig, W.; Müller, D. J.; Rog, T.; Vattulainen, I. Mechanism of allosteric regulation of *β*2-adrenergic receptor by cholesterol. Elife 2016, 5, e18432.

(60) Nagy, P. I.; Erhardt, P. W. Theoretical studies of salt-bridge formation by amino acid side chains in low and medium polarity environments. The Journal of Physical Chemistry B 2010, 114, 16436–16442.

(61) Dey, M.; Cao, C.; Sicheri, F.; Dever, T. E. Conserved intermolecular salt bridge required for activation of protein kinases PKR, GCN2, and PERK. Journal of Biological Chemistry 2007, 282, 6653–6660.

(62) Maciejko, J.; Mehler, M.; Kaur, J.; Lieblein, T.; Morgner, N.; Ouari, O.; Tordo, P.; Becker-Baldus, J.; Glaubitz, C. Visualizing specific cross-protomer interactions in the homo-oligomeric membrane protein proteorhodopsin by dynamic-nuclear-polarization-enhanced solid-state NMR. Journal of the American Chemical Society 2015, 137, 9032– 9043.

(63) Janovick, J. A.; Conn, P. M. Salt bridge integrates GPCR activation with protein trafficking. Proceedings of the National Academy of Sciences 2010, 107, 4454–4458.

(64) Guo, C.; Yang, L.; Liu, Z.; Liu, D.; Wüthrich, K. Two-dimensional NMR spectroscopy of the G protein-coupled receptor A2AAR in lipid nanodiscs. Molecules 2023, 28, 5419.

(65) Mahalingam, M.; Martínez-Mayorga, K.; Brown, M. F.; Vogel, R. Two protonation switches control rhodopsin activation in membranes. Proceedings of the National Academy of Sciences 2008, 105, 17795–17800.

(66) Ahn, K. H.; Scott, C. E.; Abrol, R.; Goddard III, W. A.; Kendall, D. A. Computationally-predicted CB1 cannabinoid receptor mutants show distinct patterns of salt-bridges that correlate with their level of constitutive activity reflected in G protein coupling levels, thermal stability, and ligand binding. Proteins: Structure, Function, and Bioinformatics 2013, 81, 1304–1317.

(67) Patra, S. M.; Chakraborty, S.; Shahane, G.; Prasanna, X.; Sengupta, D.; Maiti, P. K.; Chattopadhyay, A. Differential dynamics of the serotonin1A receptor in membrane bilayers of varying cholesterol content revealed by all atom molecular dynamics simulation. Molecular membrane biology 2015, 32, 127–137.

(68) Khelashvili, G.; Grossfield, A.; Feller, S. E.; Pitman, M.; Weinstein, H. Structural and dynamic effects of cholesterol at preferred sites of interaction with rhodopsin identified from microsecond length molecular dynamics simulations. Biophysical Journal 2009, 96, 678a.

(69) van Aalst, E.; Wylie, B. Cholesterol is a dose-dependent positive allosteric modulator of CCR3 ligand affinity and G protein coupling. Front Mol Biosci 8: 724603. 2021.

(70) Kimura, T.; Yeliseev, A. A.; Vukoti, K.; Rhodes, S. D.; Cheng, K.; Rice, K. C.; Gawrisch, K. Recombinant cannabinoid type 2 receptor in liposome model activates g protein in response to anionic lipid constituents. Journal of Biological Chemistry 2012, 287, 4076–4087.

(71) Gorvin, C. M.; Babinsky, V. N.; Malinauskas, T.; Nissen, P. H.; Schou, A. J.; Hanyaloglu, A. C.; Siebold, C.; Jones, E. Y.; Hannan, F. M.; Thakker, R. V. A calcium-sensing receptor mutation causing hypocalcemia disrupts a transmembrane salt bridge to activate *β*-arrestin–biased signaling. Science signaling 2018, 11, eaan3714.

(72) Hilger, D.; Kumar, K. K.; Hu, H.; Pedersen, M. F.; O’Brien, E. S.; Giehm, L.; Jennings, C.; Eskici, G.; Inoue, A.; Lerch, M. et al. Structural insights into differences in G protein activation by family A and family B GPCRs. Science 2020, 369, eaba3373.

(73) Janovick, J. A.; Pogozheva, I. D.; Mosberg, H. I.; Conn, P. M. Salt bridges overlapping the gonadotropin-releasing hormone receptor agonist binding site reveal a coincidence detector for G protein-coupled receptor activation. The Journal of Pharmacology and Experimental Therapeutics 2011, 338, 430–442.

(74) Boerner, T. J.; Deems, S.; Furlani, T. R.; Knuth, S. L.; Towns, J. Practice and Experience in Advanced Research Computing 2023: Computing for the Common Good; 2023; pp 173–176.

