## Supporting Information for "Lipid-Mediated Modulation of mGluR2 Embedded in Micelle and Bilayer Environments: Insights from Molecular Dynamics"

### List of Figures


S1 Time series of C$\alpha$ RMSD for mGluR2 embedded in a micelle . . . . . . . . . S3

S2 RMSD analysis of mGluR2 protomer stability in the inactive state . . . . . . S4

S3 RMSD analysis of mGluR2 protomer stability in the active state . . . . . . . S5

S4 Interprotomer distance analysis of mGluR2 in micelle systems . . . . . . . . S6

S5 Interprotomer angle analysis of mGluR2 in micelle systems . . . . . . . . . . S7

S6 C$\alpha$ RMSD analysis of individual helices in mGluR2 (inactive state, micelle) . S8

S7 C$\alpha$ RMSD analysis of individual helices in mGluR2 (active state, micelle) . . S9

S8 Time series of C$\alpha$ RMSD for mGluR2 embedded in a bilayer . . . . . . . . . S10

S9 RMSD analysis of mGluR2 protomers embedded in a bilayer (inactive state) S11

S10 RMSD analysis of mGluR2 protomers embedded in a bilayer (active state) . S12

S11 Interprotomer distance analysis of mGluR2 embedded in a bilayer . . . . . . S13

S12 Interprotomer angle analysis of mGluR2 embedded in a bilayer . . . . . . . . S14

S13 C$\alpha$ RMSD analysis of individual helices of mGluR2 embedded in a bilayer (inactive state) . . . . . . . . . . . . . . . . . . . . . . . . . . . . . . . . . . S15

S14 C$\alpha$ RMSD analysis of individual helices of mGluR2 embedded in a bilayer (active state) . . . . . . . . . . . . . . . . . . . . . . . . . . . . . . . . . . . S16

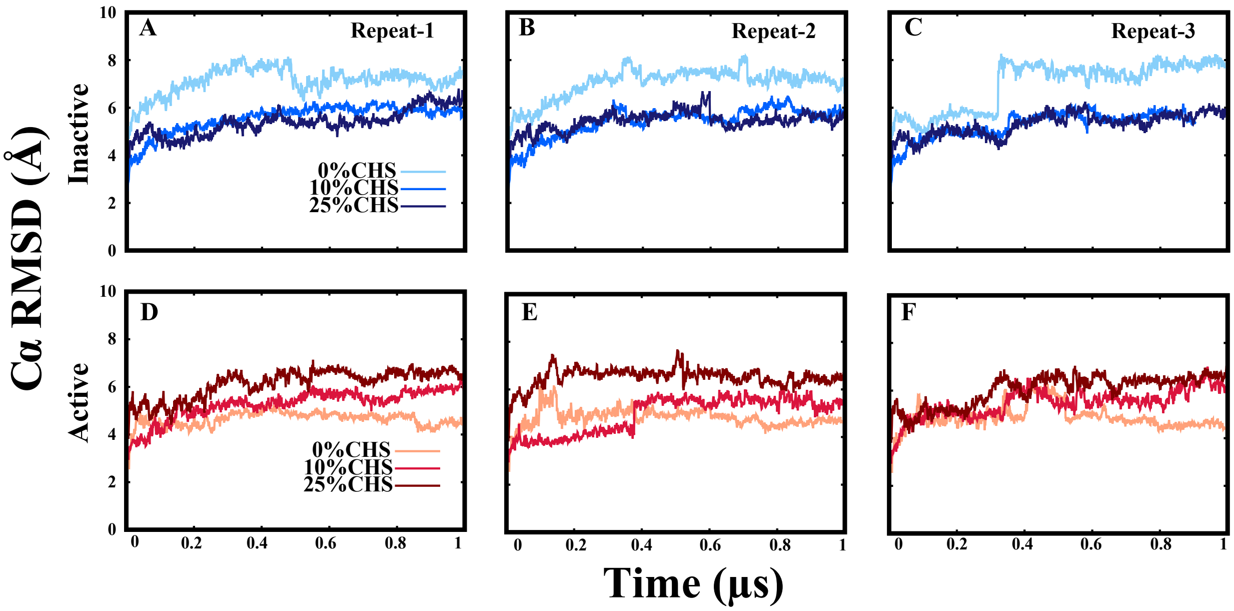


**Fig. S1. Time series of C$\alpha$ RMSD for mGluR2 embedded in a micelle.** Time evolution of the C$\alpha$ RMSD for the full mGluR2 receptor in the inactive (A–C) and active (D–F) states, embedded in a micelle, under varying CHS concentrations (0%, 10%, and 25%). Each panel represents one of three independent 1 $\mu$s simulation repeats. The results illustrate CHS-dependent stabilization in the inactive state and increased flexibility in the active state, particularly at higher CHS concentrations.

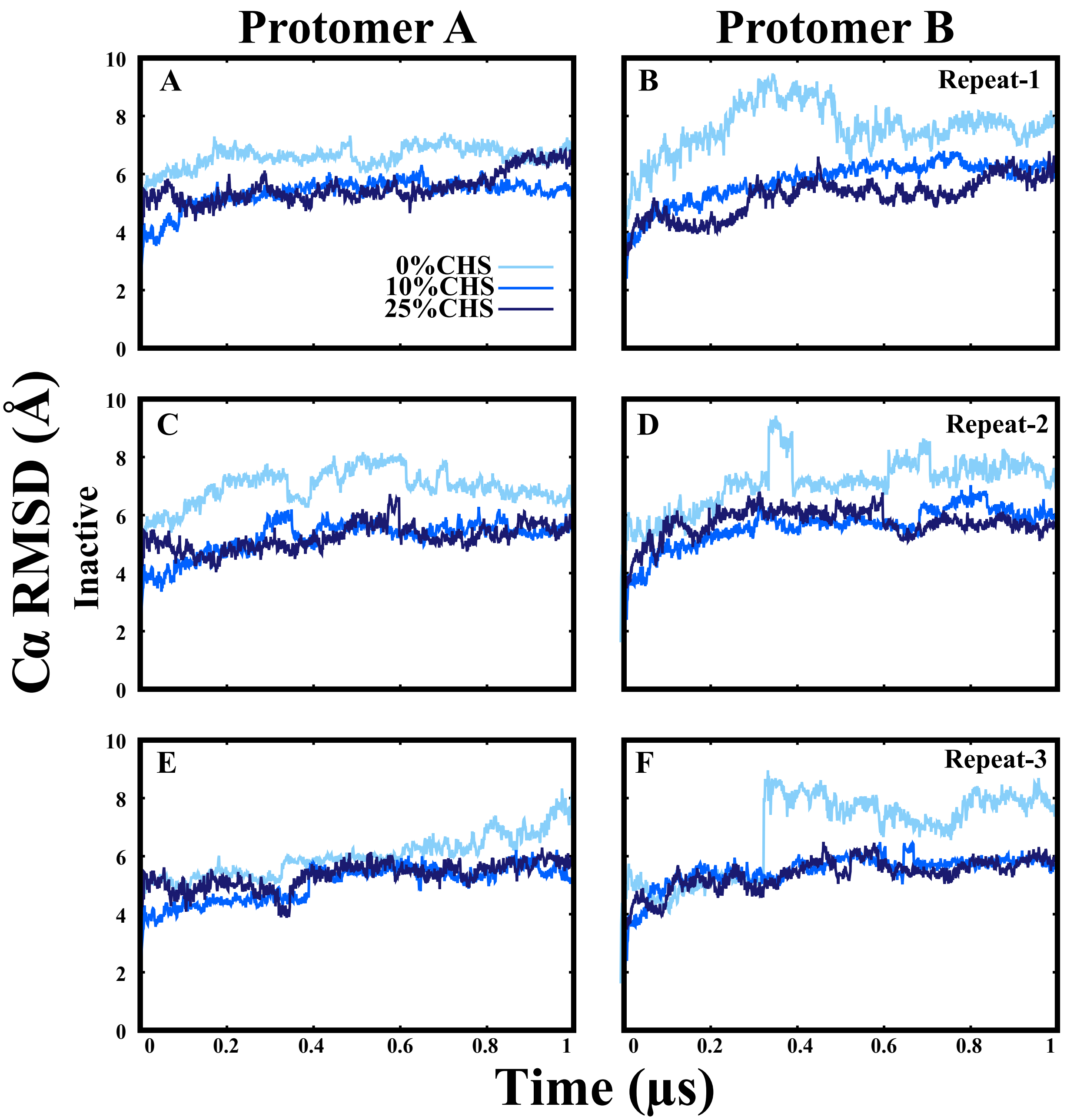


**Fig. S2. RMSD analysis of mGluR2 protomer stability in the inactive state.** Time series of C$\alpha$ RMSD for protomer A (panels A, C, E) and protomer B (panels B, D, F) of the mGluR2 receptor in the inactive state, embedded in a micelle, under varying CHS concentrations (0%, 10%, and 25%). Each row corresponds to a distinct simulation repeat (Repeat-1: A–B; Repeat-2: C–D; Repeat-3: E–F). Simulations were conducted for 1 $\mu$s to evaluate the structural stability of individual protomers under cholesterol variation. The results demonstrate CHS-dependent trends in conformational dynamics over time.

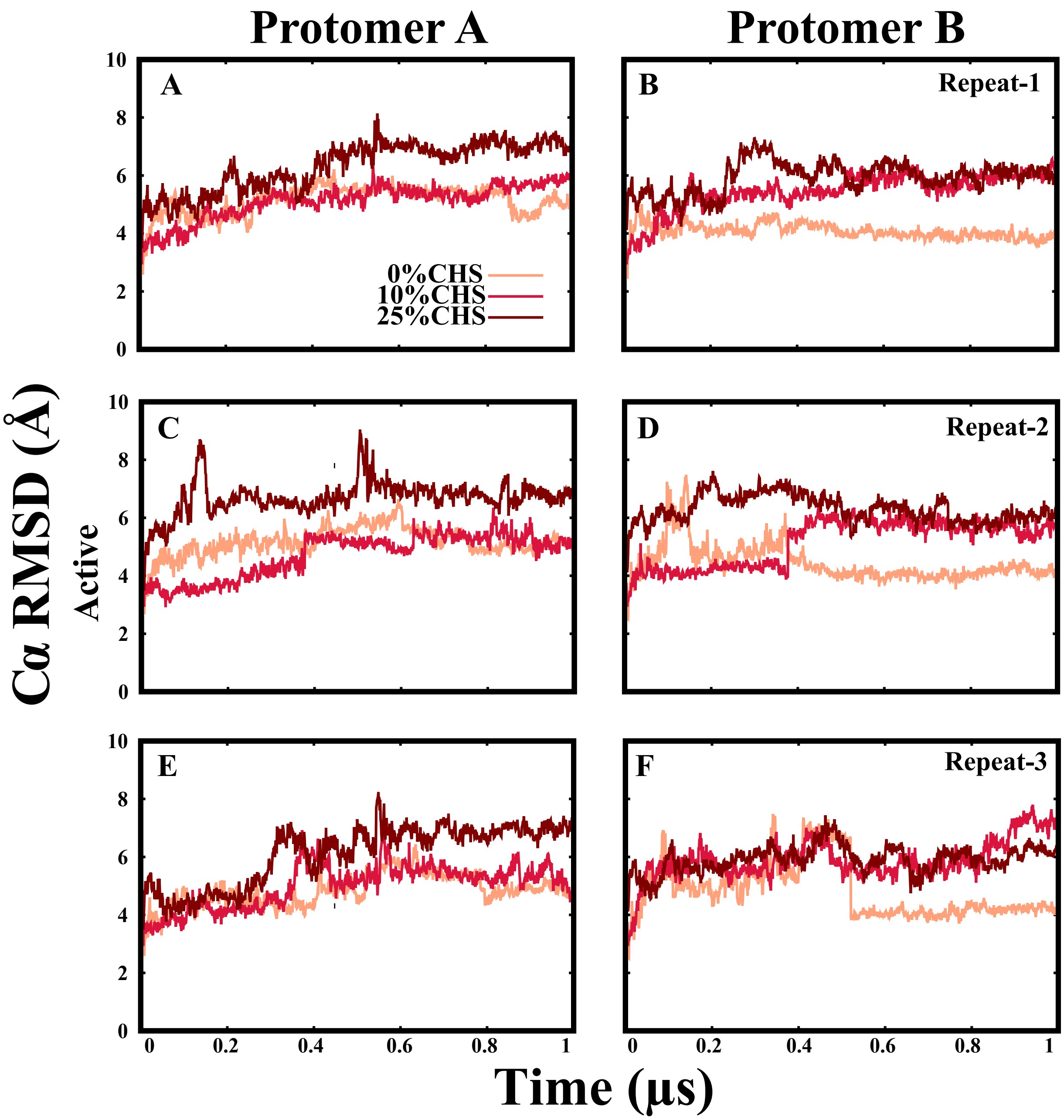


**Fig. S3. RMSD analysis of mGluR2 protomer stability in the active state.** Time series of C$\alpha$ RMSD for protomer A (panels A, C, E) and protomer B (panels B, D, F) of the mGluR2 receptor in the active state, embedded in a micelle, under varying CHS concentrations (0%, 10%, and 25%). Each row represents one of three independent simulation repeats (Repeat-1: A–B; Repeat-2: C–D; Repeat-3: E–F). Simulations were performed for 1 $\mu$s to assess the structural dynamics of individual protomers under different cholesterol conditions. The plots reveal CHS-dependent variations in flexibility and stability, particularly at higher concentrations.

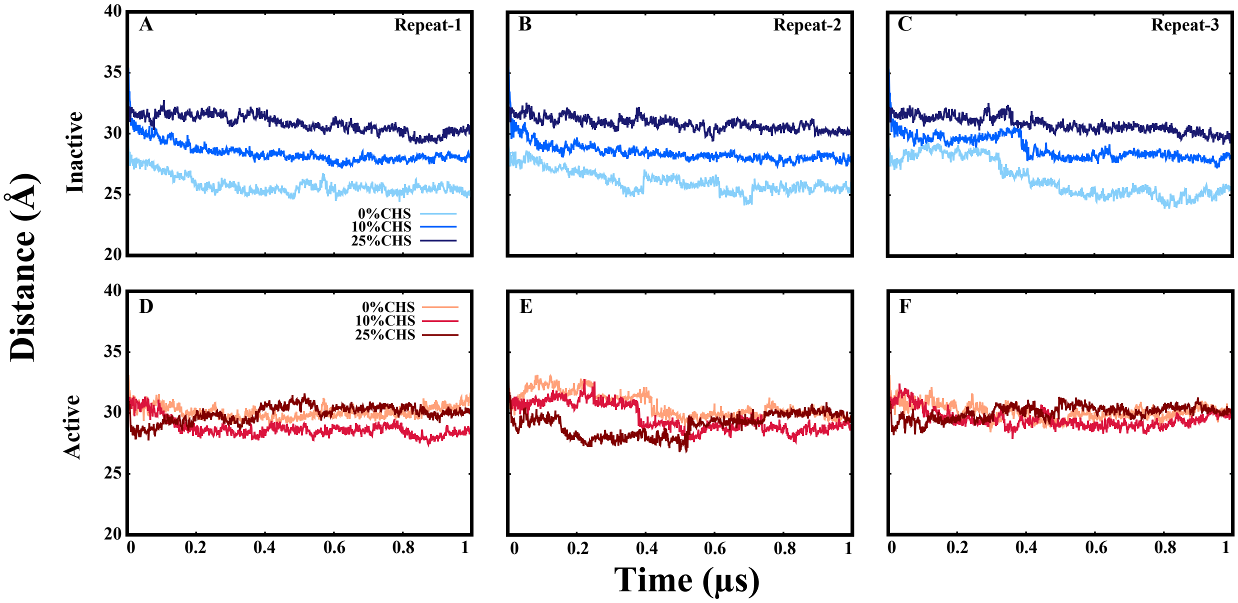


**Fig. S4. Interprotomer distance analysis of mGluR2 in micelle systems.** Time series of the interprotomer Cα–Cα distance between protomer A and protomer B of the mGluR2 receptor under varying CHS concentrations (0%, 10%, and 25%) in a micelle environment. Panels A–C represent the inactive state across three independent simulation repeats; panels D–F show the active state across the same repeats. Each simulation was run for 1 $\mu$s. The results reveal CHS-dependent modulation of the relative protomer positioning, with distinct trends between active and inactive states.

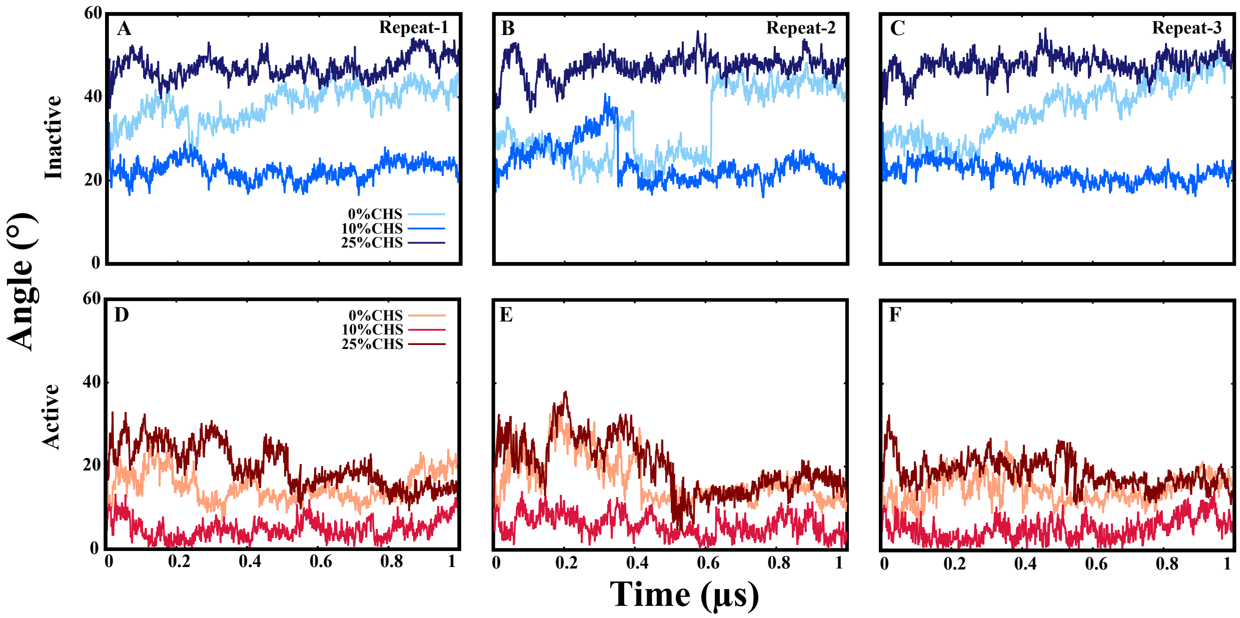


**Fig. S5. Interprotomer angle analysis of mGluR2 in micelle systems.** Time series of the angle between protomer A and protomer B of the mGluR2 receptor under varying CHS concentrations (0%, 10%, and 25%) in a micelle environment. Panels A–C represent the inactive state across three independent simulation repeats; panels D–F correspond to the active state. Each simulation was conducted for 1 $\mu$s. The results demonstrate CHS-dependent modulation of the relative protomer orientation, with more pronounced angular fluctuations observed in the inactive state compared to the active state.

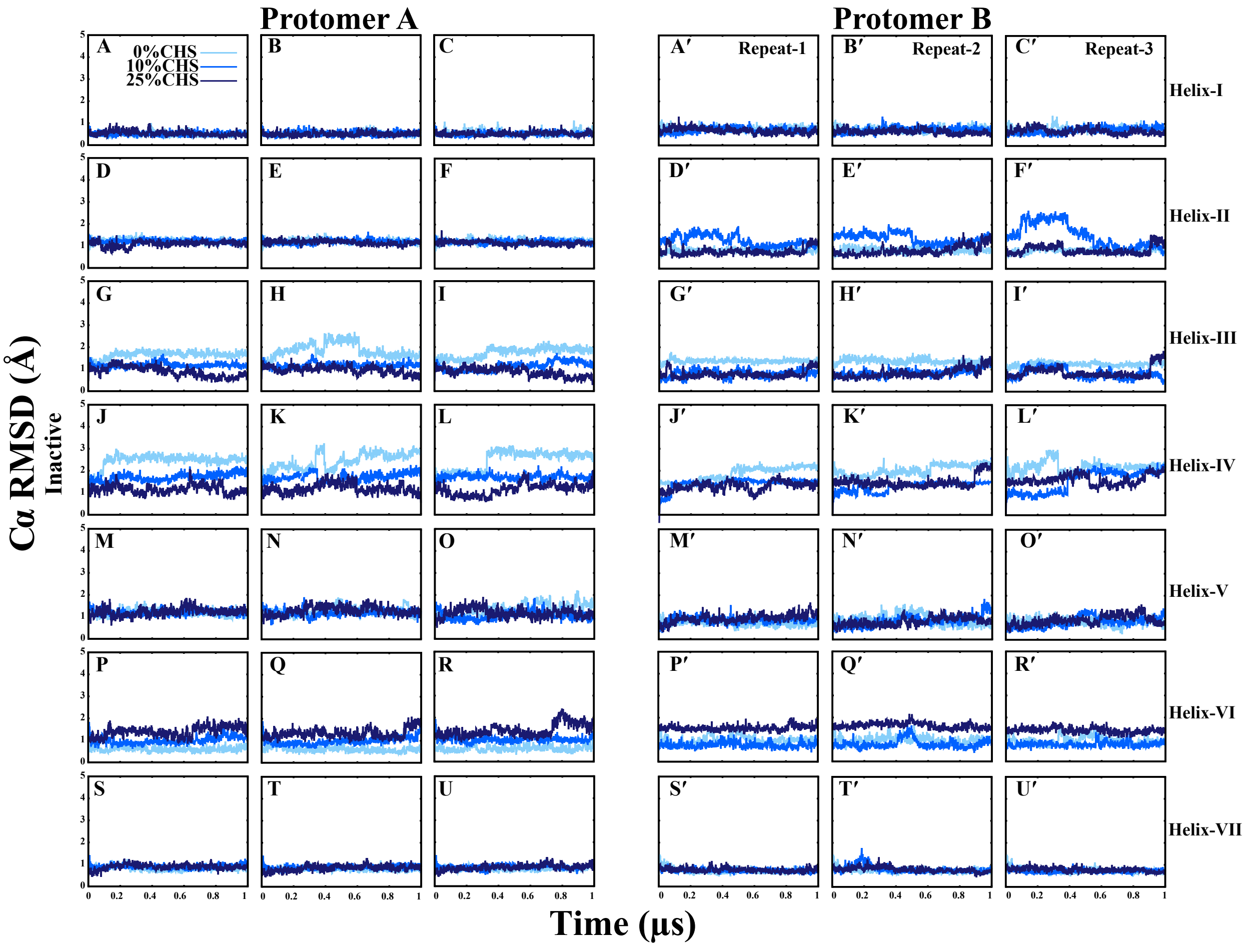


**Fig. S6. C$\alpha$ RMSD analysis of individual helices in mGluR2 (inactive state, micelle).** Time series of C$\alpha$ RMSD for transmembrane helices I–VII in protomer A (panels A–U) and protomer B (panels A′–U′) of the mGluR2 receptor in the inactive state, embedded in a micelle, at CHS concentrations of 0%, 10%, and 25%. Each panel displays RMSD fluctuations for a specific helix across three independent 1 $\mu$s simulation repeats. Helices are arranged in rows: helix I (A–C, A′–C′), helix II (D–F, D′–F′), helix III (G–I, G′–I′), helix IV (J–L, J′–L′), helix V (M–O, M′–O′), helix VI (P–R, P′–R′), and helix VII (S–U, S′–U′). The results highlight CHS-dependent effects on the stability and flexibility of individual transmembrane helices.

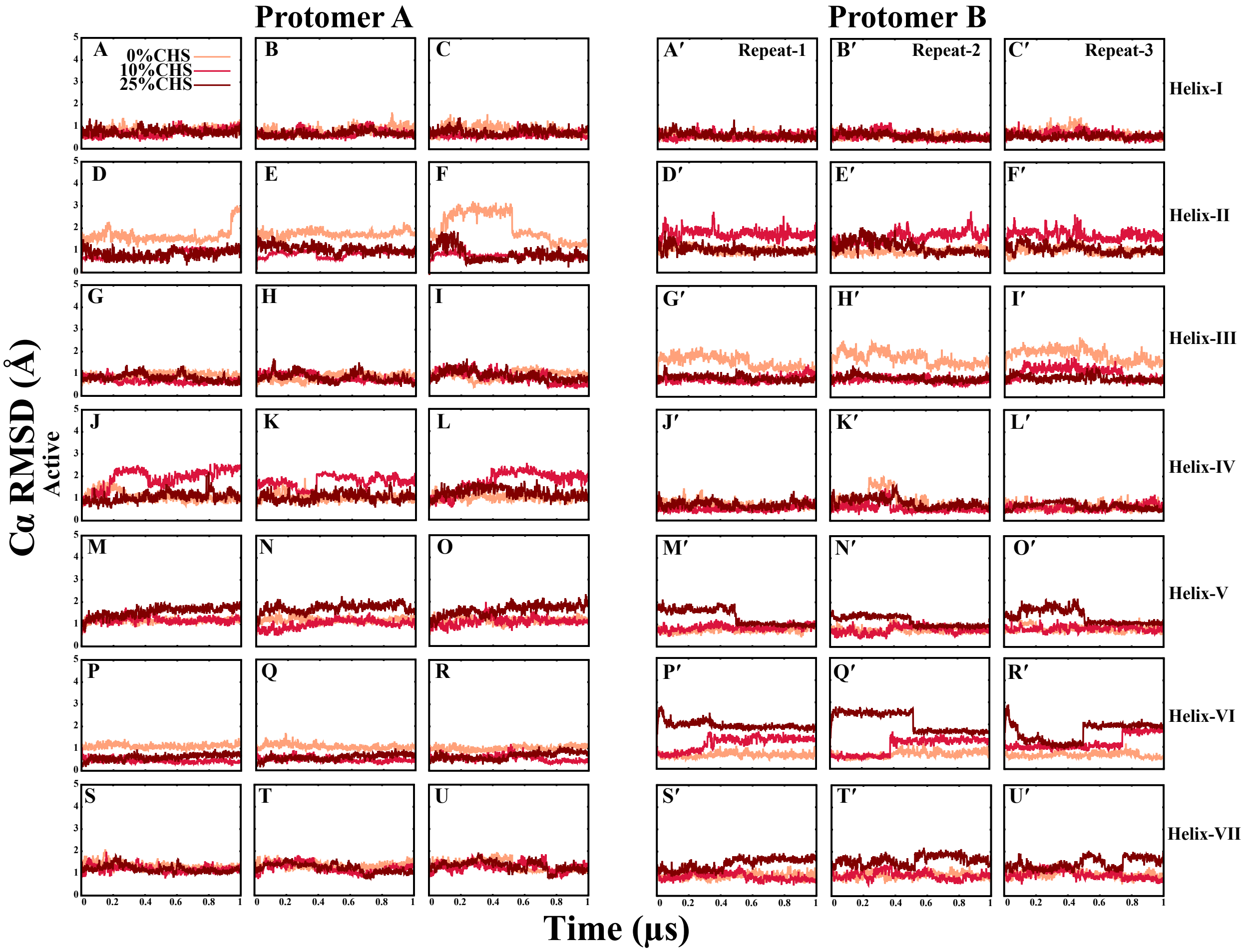


**Fig. S7. Cα RMSD analysis of individual helices in mGluR2 (active state, micelle).** Time series of Cα RMSD for transmembrane helices I–VII in protomer A (panels A–U) and protomer B (panels A′–U′) of the mGluR2 receptor in the active state, embedded in a micelle, at CHS concentrations of 0%, 10%, and 25%. Each panel displays RMSD fluctuations for a specific helix across three independent 1 $\mu$s simulation repeats. Helices are arranged in rows: helix I (A–C, A′–C′), helix II (D–F, D′–F′), helix III (G–I, G′–I′), helix IV (J–L, J′–L′), helix V (M–O, M′–O′), helix VI (P–R, P′–R′), and helix VII (S–U, S′–U′). The results highlight CHS-dependent effects on the structural flexibility and stability of individual transmembrane helices.

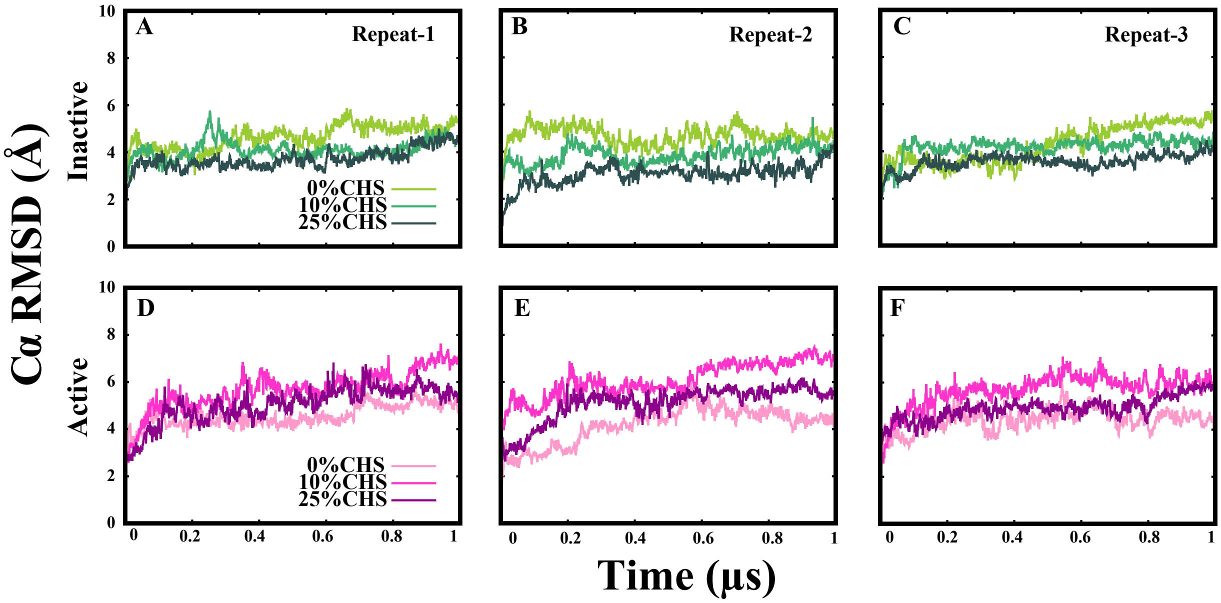


**Fig. S8. Time series of Cα RMSD for mGluR2 embedded in a bilayer.** Time evolution of the Cα RMSD for the mGluR2 receptor in the inactive (A–C) and active (D–F) states, embedded in a lipid bilayer, under varying CHS concentrations (0%, 10%, and 25%). Panels A–C and D–F correspond to three independent simulation repeats for the inactive and active states, respectively. All simulations were carried out for 1 $\mu$s to assess the structural stability of the receptor in a bilayer environment with different cholesterol levels.

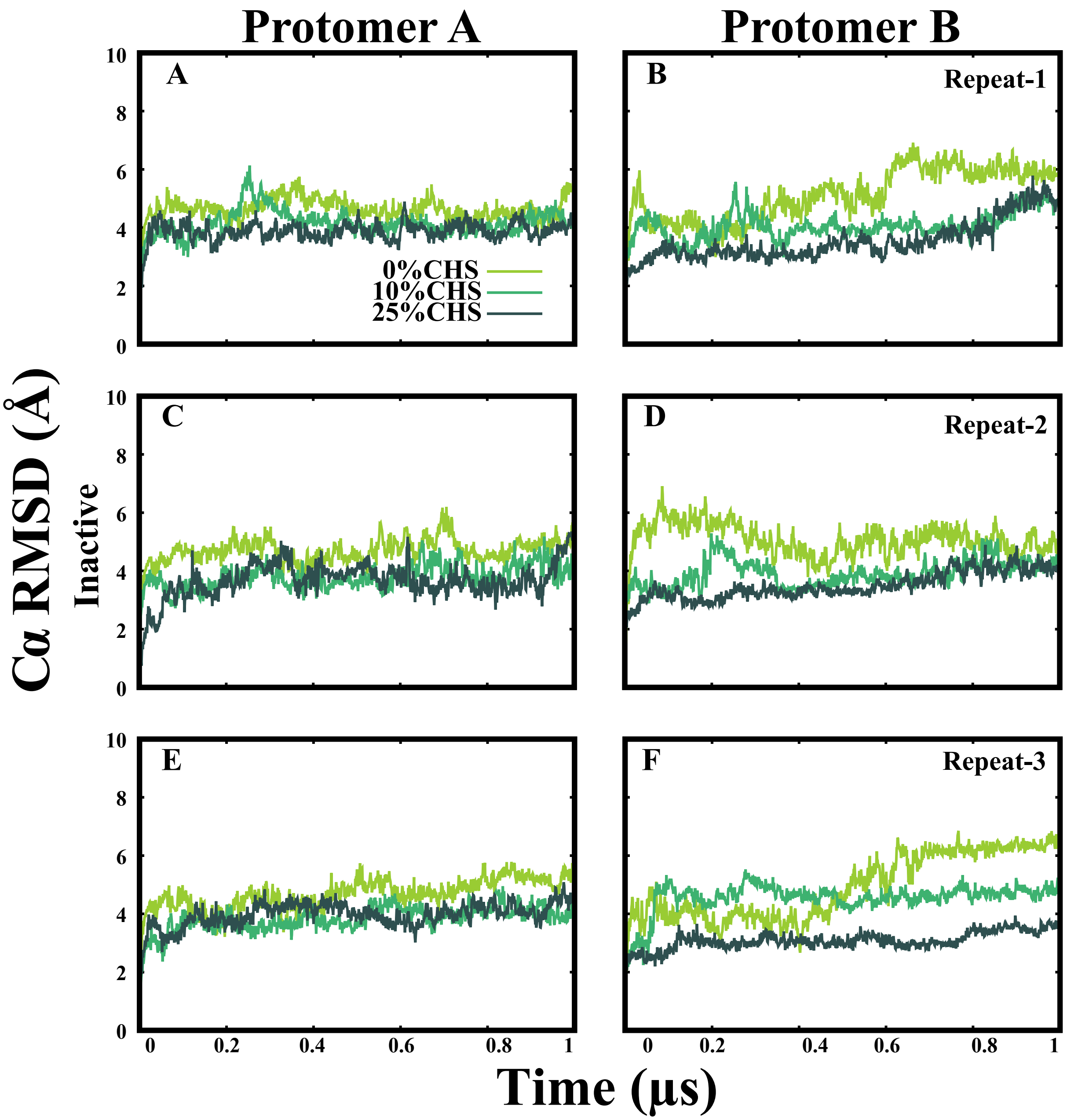


**Fig. S9. RMSD analysis of mGluR2 protomers embedded in a bilayer (inactive state).** Time series of Cα RMSD for protomer A (panels A–C) and protomer B (panels D–F) of the mGluR2 receptor in the inactive state, embedded in a lipid bilayer, under varying CHS concentrations (0%, 10%, and 25%). Each row represents one of three independent 1 $\mu$s simulation repeats. The data highlight CHS-dependent differences in protomer stability and dynamics within the bilayer environment.

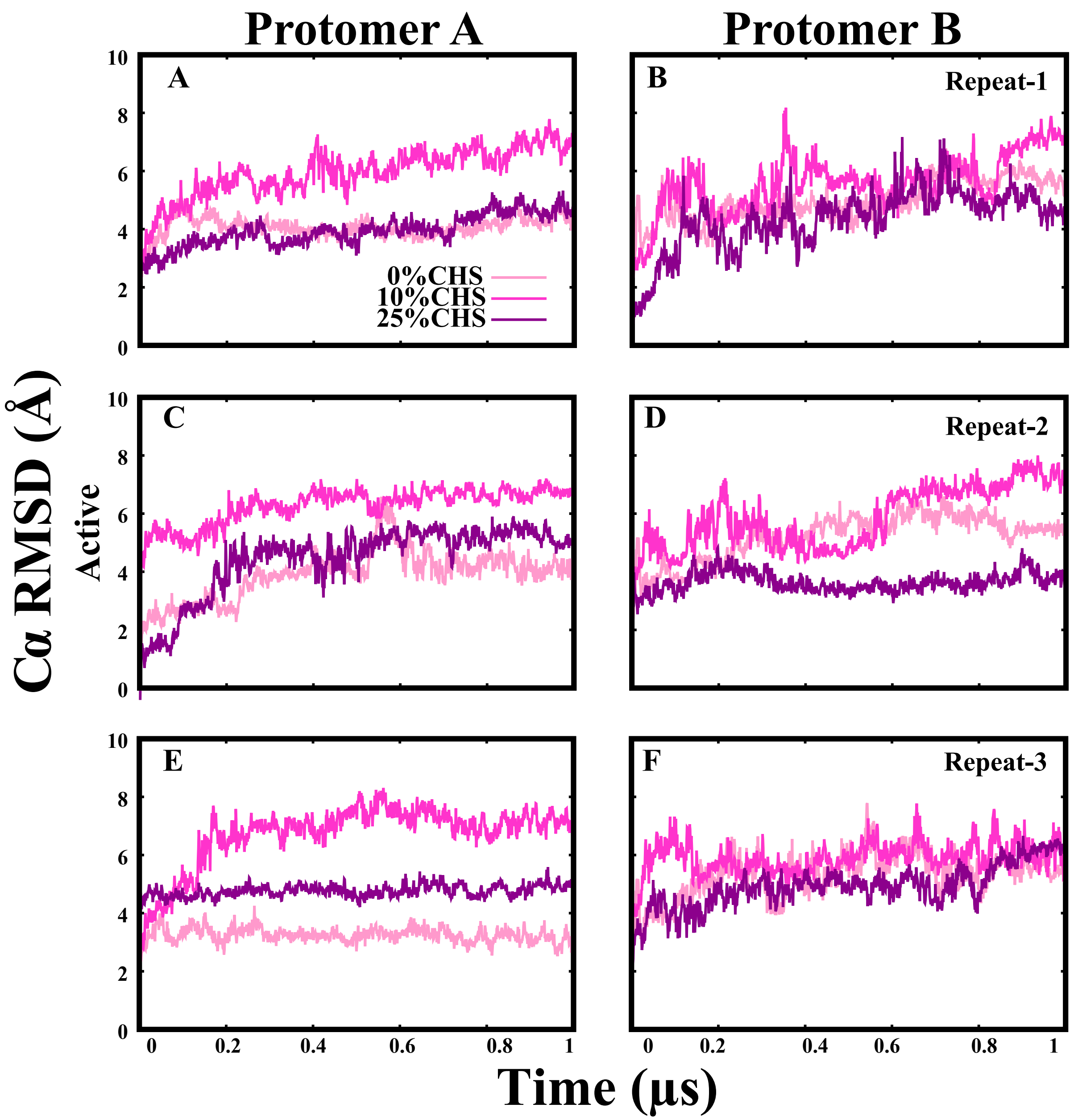


**Fig. S10. RMSD analysis of mGluR2 protomers embedded in a bilayer (active state).** Time series of Cα RMSD for protomer A (panels A–C) and protomer B (panels D–F) of the mGluR2 receptor in the active state, embedded in a lipid bilayer, under varying CHS concentrations (0%, 10%, and 25%). Each row corresponds to one of three independent 1 $\mu$s simulation replicates. The data illustrate CHS-dependent differences in conformational flexibility and stability of the receptor within the bilayer environment.

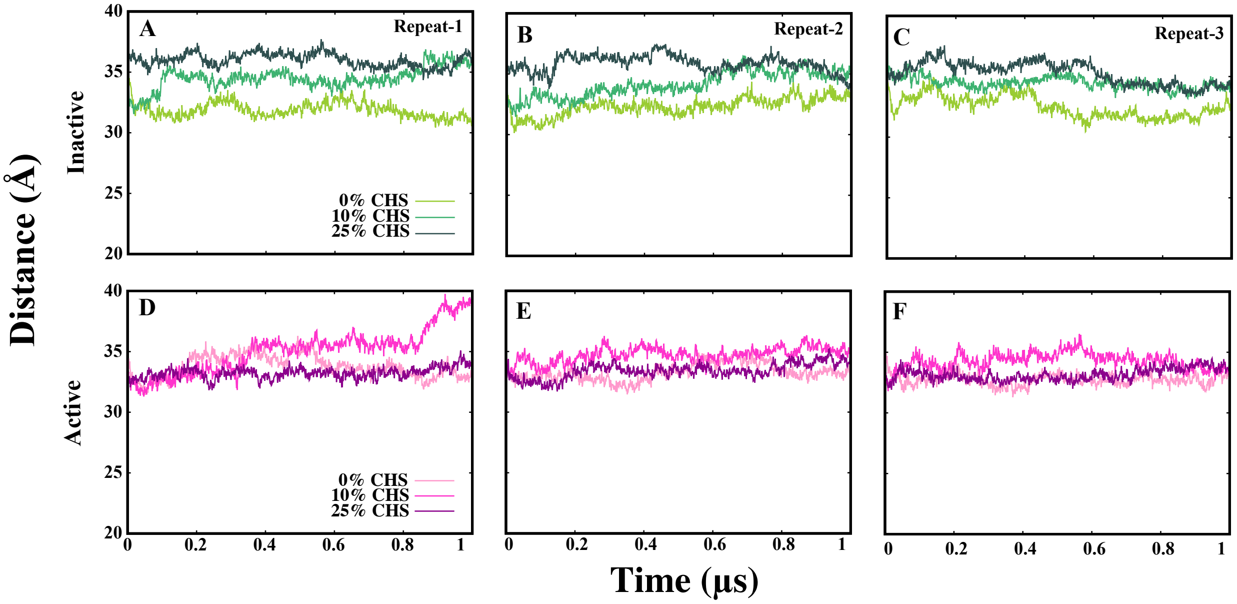


**Fig. S11. Interprotomer distance analysis of mGluR2 embedded in a bilayer.** Time series of the interprotomer C$\alpha$–C$\alpha$ distance between protomer A and protomer B of the mGluR2 receptor under varying CHS concentrations (0%, 10%, and 25%) in a lipid bilayer environment. Panels A–C correspond to the inactive state across three independent simulation replicates, while panels D–F represent the active state. Each simulation was conducted for 1 $\mu$s. The plots highlight CHS-dependent changes in protomer separation, indicating dynamic modulation of dimeric stability in different conformational states.

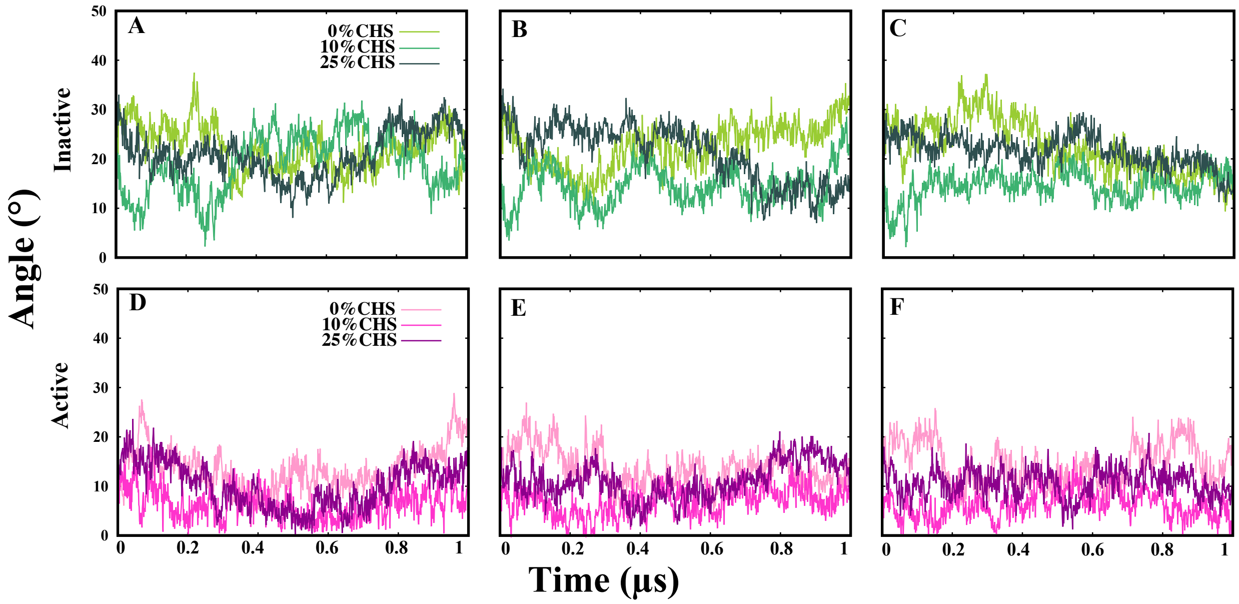


**Fig. S12. Interprotomer angle analysis of mGluR2 embedded in a bilayer.** Time series of the interprotomer angle between protomer A and protomer B of the mGluR2 receptor under varying CHS concentrations (0%, 10%, and 25%) in a lipid bilayer environment. Panels A–C correspond to the inactive state across three independent simulation replicates, while panels D–F represent the active state. Each simulation was run for 1 $\mu$s. The results illustrate CHS-dependent modulation of the angular relationship between protomers, revealing greater flexibility in the active state and more defined angular positioning in the inactive conformation.

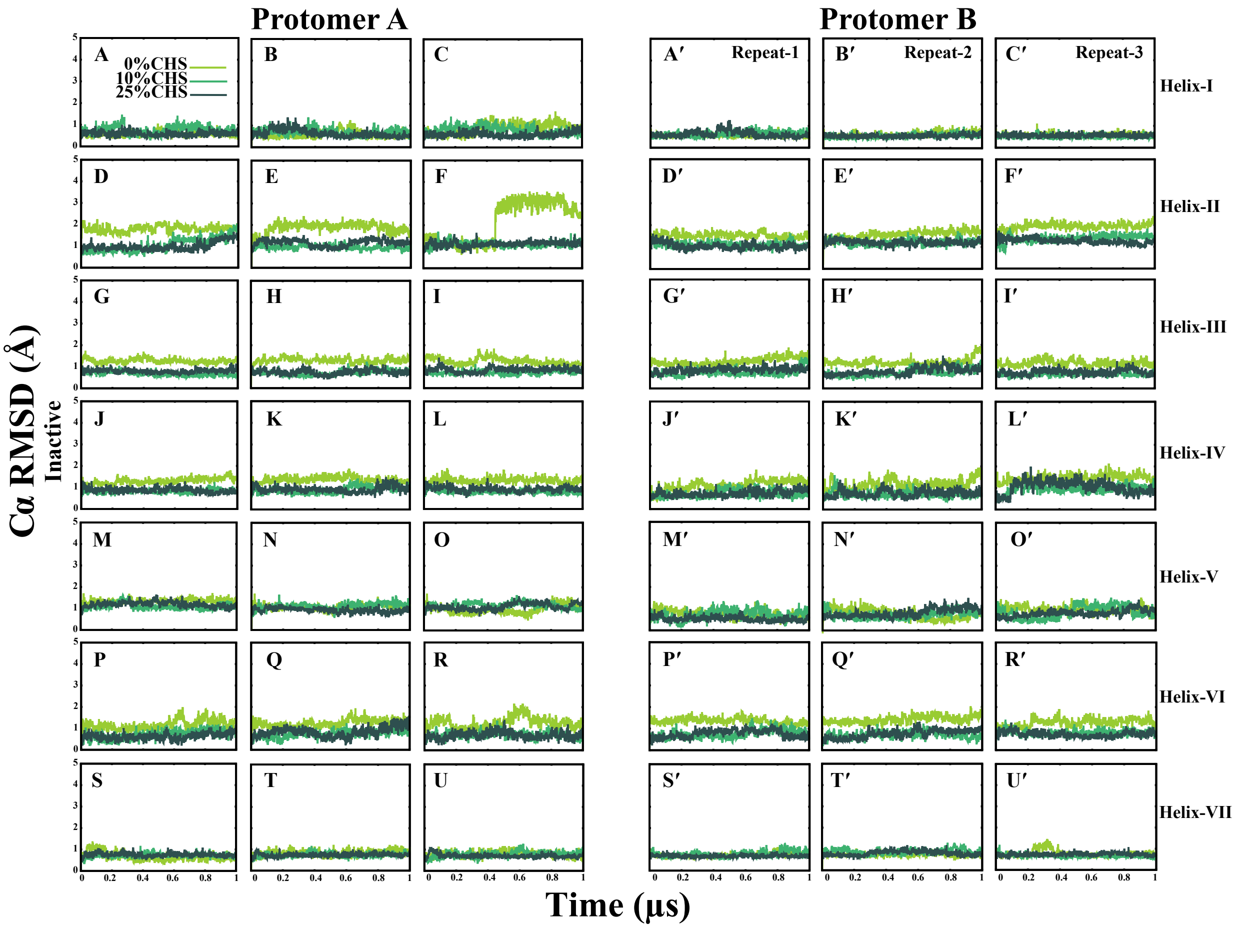


**Fig. S13. Cα RMSD analysis of individual helices of mGluR2 embedded in a bilayer (inactive state).** Time series of Cα RMSD for individual transmembrane helices (I–VII) in protomer A (A–U) and protomer B (A′–U′) of the mGluR2 receptor in the inactive state, embedded in a lipid bilayer, at CHS concentrations of 0%, 10%, and 25%. Each panel displays RMSD fluctuations for a specific helix across three independent simulation repeats. Helices are arranged by rows: Helix I (A–C, A′–C′), Helix II (D–F, D′–F′), Helix III (G–I, G′–I′), Helix IV (J–L, J′–L′), Helix V (M–O, M′–O′), Helix VI (P–R, P′–R′), and Helix VII (S–U, S′–U′). Simulations were performed for 1 $\mu$s to assess how CHS modulates the flexibility and conformational stability of individual helices within the bilayer environment.

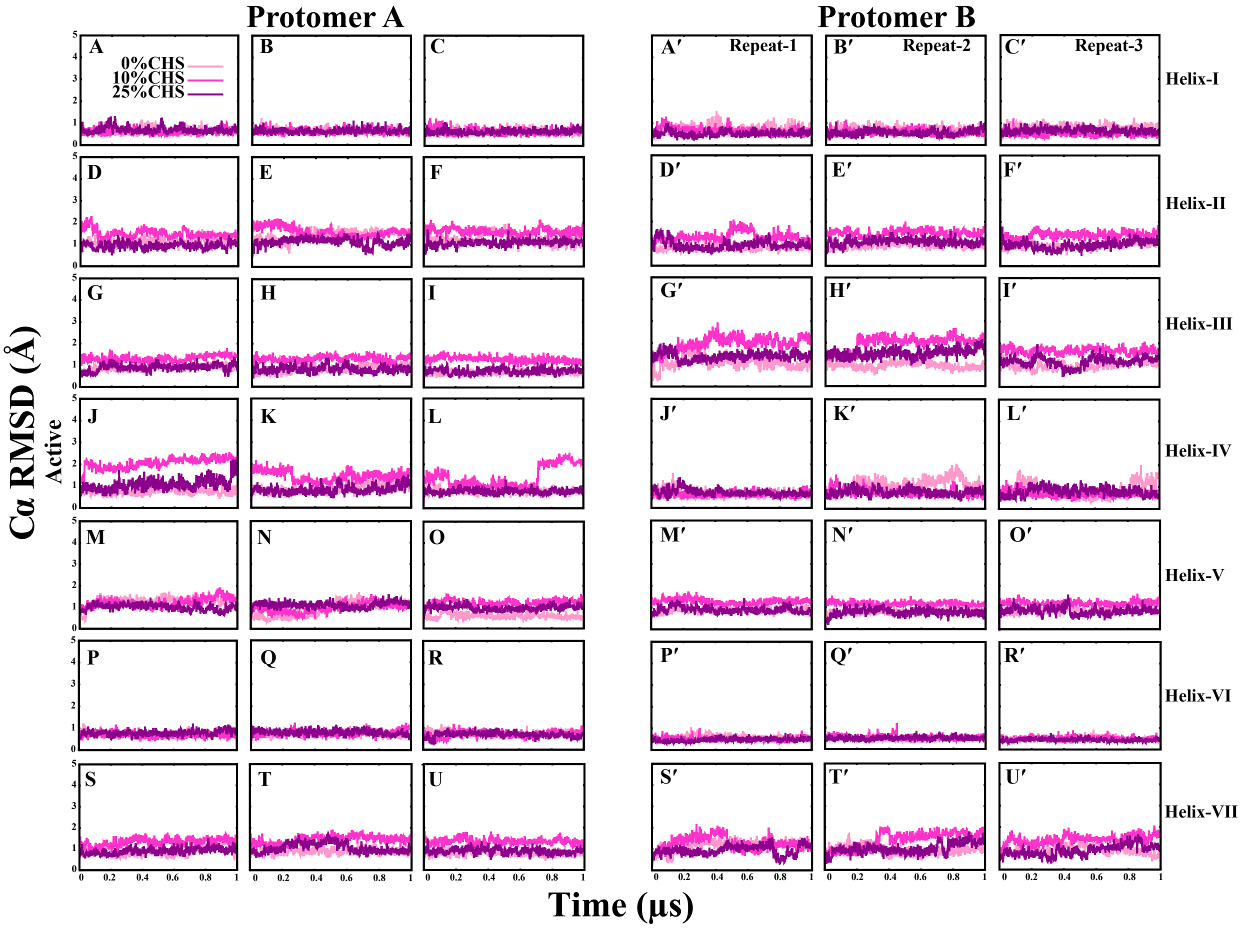


**Fig. S14. Cα RMSD analysis of individual helices of mGluR2 embedded in a bilayer (active state).** Time series of Cα RMSD for individual transmembrane helices (I–VII) in protomer A (A–U) and protomer B (A′–U′) of the mGluR2 receptor in the active state, embedded in a lipid bilayer, at CHS concentrations of 0%, 10%, and 25%. Each panel displays RMSD fluctuations for a specific helix across three independent simulation repeats. Helices are arranged by rows: Helix I (A–C, A′–C′), Helix II (D–F, D′–F′), Helix III (G–I, G′–I′), Helix IV (J–L, J′–L′), Helix V (M–O, M′–O′), Helix VI (P–R, P′–R′), and Helix VII (S–U, S′–U′). Simulations were performed for 1 $\mu$s to examine the CHS-dependent effects on the flexibility and structural stability of individual helices within the bilayer environment.
